# A pan-genomic and methylomic analysis reveals a distinct signature in *Vibrio alginolyticus* isolated from wild fish

**DOI:** 10.64898/2026.09.10.750592

**Authors:** Liang Zhong, Yuxuan Zhang, Meng Yan, Runsheng Li, Wenlong Cai

## Abstract

*Vibrio alginolyticus* is a ubiquitous opportunistic pathogen in estuarine and marine ecosystems and a leading cause of vibriosis in humans and aquatic animals. Yet genomic and epigenomic landscapes of *V. alginolyticus* from wild fish remain poorly defined. Here, we present a pan-genomic and methylomic framework based on 89 high-quality *V. alginolyticus* genomes, including 46 newly sequenced isolates recovered from 105 wild marine fish representing 17 species in Hong Kong waters of the South China Sea, plus all publicly available complete genomes. We reported a 32.38% prevalence of *V. alginolyticus* in wild fish. Phylogenomic analysis resolved four distinct clades, with Clade IV dominated by wild-fish isolates and characterized by low virulence and low antimicrobial resistance. Pan-genomic analysis revealed a closed pan-genome with a substantially depleted accessory genome in Clade IV. We identified a total of 763 antimicrobial resistance (AMR) genes from 89 strains, which covered 32 gene types and spanned four resistance mechanisms, with efflux pumps as the most prevalent strategy. Critically, resistance genes were almost exclusively chromosomal rather than plasmid-borne. Virulence profiling confirmed the presence of *tlh* and T6SS genes but the absence of the high-risk human pathogenic factors *tdh* and *ctxB*. Methylomic analysis using nanopore sequencing uncovered 355 DNA methyltransferases and 140 strain-specific methylation motifs with dominance by 6mA. Notably, 71.90% of methyltransferases resided in the accessory genome and were disseminated by mobile genetic elements, especially plasmids. Motif combinations were highly strain-specific and largely decoupled from phylogeny, except for a shared motif signature defining Clade IV. The GATC motif was essential across all *V. alginolyticus* strains and showed significant enrichment in virulence gene regions but not in AMR gene loci, revealing differentiated epigenetic modification characteristics. This study nearly doubled the number of high-quality complete genomes available for *V. alginolyticus* and provides the first comprehensive methylomic characterization for this species. Our findings reveal a phylogenetically distinct, low-virulence, low-resistance *V. alginolyticus* clade widely shared among wild fish, with important implications for One Health surveillance and evolutionary adaptation of marine pathogens.

## 1 Introduction

*Vibrio* species are curved Gram-negative bacilli widely distributed in rivers, estuaries, and coastal oceans (de Souza Valente and Wan, 2021). Among more than 130 described species, 12 species are pathogenic to humans, including *V. cholerae*, *V. parahaemolyticus*, *V. vulnificus*, and *V. alginolyticus* (Ceccarelli et al., 2019). As a dominant marine *Vibrio* species, *V. alginolyticus* causes severe disease and mass mortality in farmed fish, shrimp, and shellfish, resulting in substantial economic losses globally (Liu et al., 2004; Mohamad et al., 2019; Wang et al., 2016). It is also the second most common *Vibrio* species associated with human infections (CDC, 2012), causing soft-tissue infections, sepsis, and gastroenteritis following contact with contaminated water or consumption of raw seafood (Altekruse et al., 2000; Hlady and Klontz, 1996; Jacobs Slifka et al., 2017; Sganga et al., 2009; Uh et al., 2001). Despite its clinical, veterinary, and ecological importance, large-scale genomic and epigenomic data for *V. alginolyticus* from wild fish remain scarce, limiting understanding of its prevalence, virulence, antimicrobial resistance (AMR), and evolution in natural marine ecosystems.

DNA methylation is regarded as a significant epigenetic regulator in prokaryotes. Bacterial DNA methylation was formerly considered a by-product of phage resistance mechanisms associated with restriction-modification (RM) systems (Seong et al., 2021). However, recent studies have demonstrated that it is crucial not only for regulating the host defense system but also for modulating the cell cycle, gene expression, and virulence (Seong et al., 2021; Tisza et al., 2023). DNA methylation encompasses three forms: C^5^-methyl-cytosine (5mC), N^4^-methyl-cytosine (4mC), and N^6^-methyl-adenine (6mA). Among these, 5mC predominantly occurs in eukaryotes, while 6mA and 4mC are primarily present in bacteria, with 6mA being the most prevalent (Beaulaurier et al., 2019). DNA methylation is intrinsically linked to DNA methyltransferases, which are components of the RM system. Methyltransferases facilitate the methylation of DNA bases by transferring methyl groups from S-adenosylmethionine to bases within specified DNA sequences to execute necessary physiological functions (Casadesús, 2016). Long-read nanopore sequencing enables direct, genome-wide detection of modified bases, making it a powerful tool for bacterial methylomic analysis. To date, no comprehensive methylomic study has been reported for *V. alginolyticus.* Therefore, understanding DNA methylation and methyltransferases is essential to unraveling bacterial virulence and drug resistance.

In this study, we sampled 105 wild marine fish from 17 species in Hong Kong waters, isolated 46 *V. alginolyticus* strains, and generated complete circular genomes and methylomes using nanopore sequencing. By integrating these data with all public complete genomes, we constructed a high-resolution phylogeny, analyzed the pan-genome, characterized the resistome, virulome, plasmidome, and epigenome, and identified a distinct wild-fish-adapted clade with low virulence and low AMR. Our results provide a foundational resource for *V. alginolyticus* surveillance, evolutionary biology, and One Health risk assessment.

## 2 Materials and Methods

### 2.1 Fish sampling

During October and December 2022, three auxiliary vessels fitted with lighting devices were stationed at three distinct sampling sites within Tolo Harbor, Hong Kong, with specific geographic coordinates of 22.449188°N, 114.268879°E; 22.453560°N, 114.275581°E; and 22.457316°N, 114.279235°E. These lighting facilities were used for no less than 3 h to aggregate fish at each sampling site. Fish specimens at each sampling site were then collected separately using a seine net (300 m × 100 m, 1.5 cm stretched mesh) deployed from a main boat. After harvesting, professional fishermen conducted preliminary species sorting of the collected fish on-site aboard the vessel. Subsequently, individual fish samples of each confirmed species were sealed in sterile polyethylene bags and immediately stored in ice boxes, then transported to the Aquatic Animal Health Laboratory of City University of Hong Kong for subsequent analysis. A total of 105 wild marine fish specimens belonging to 17 different species were obtained (Table 1).

**Table 1.**
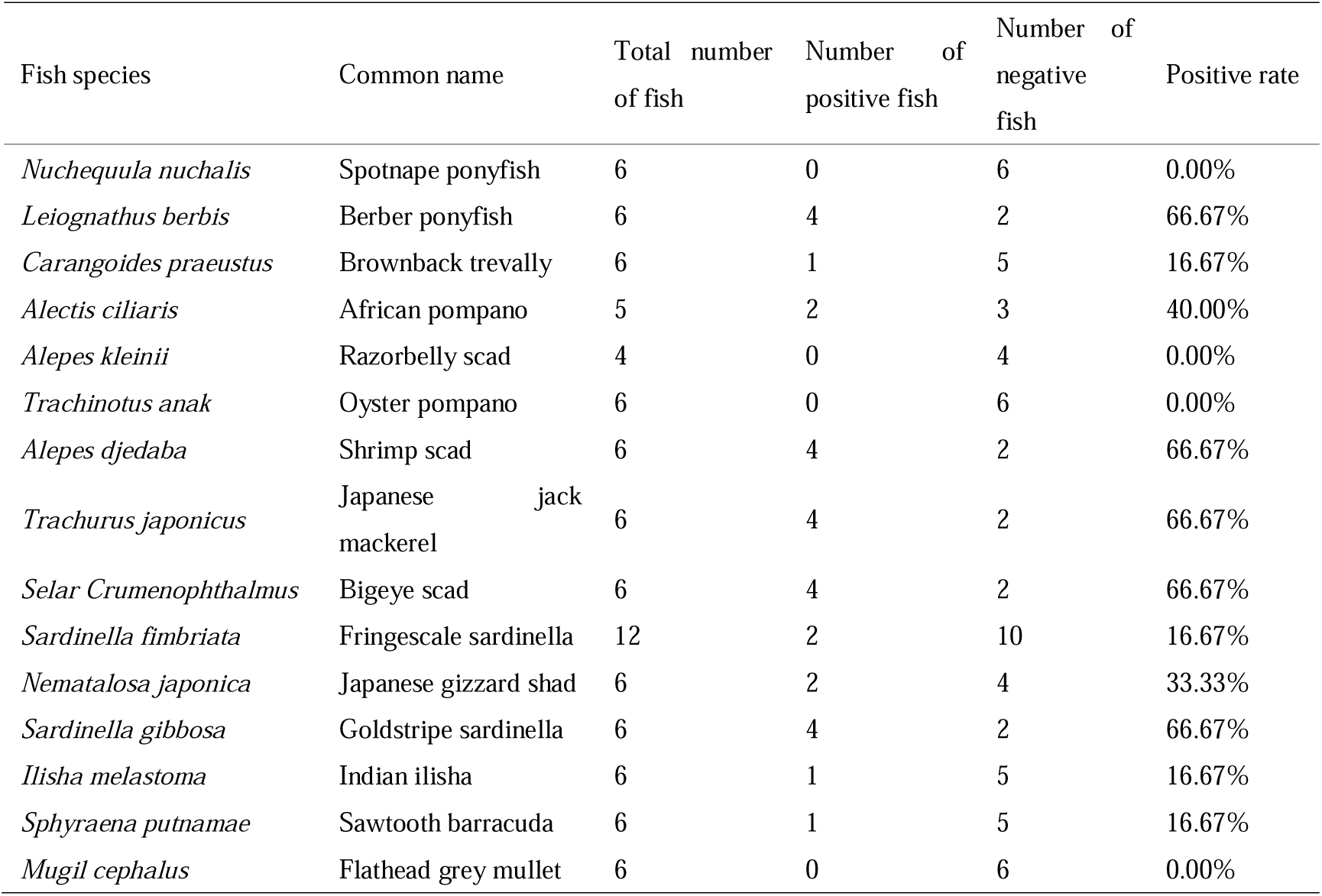
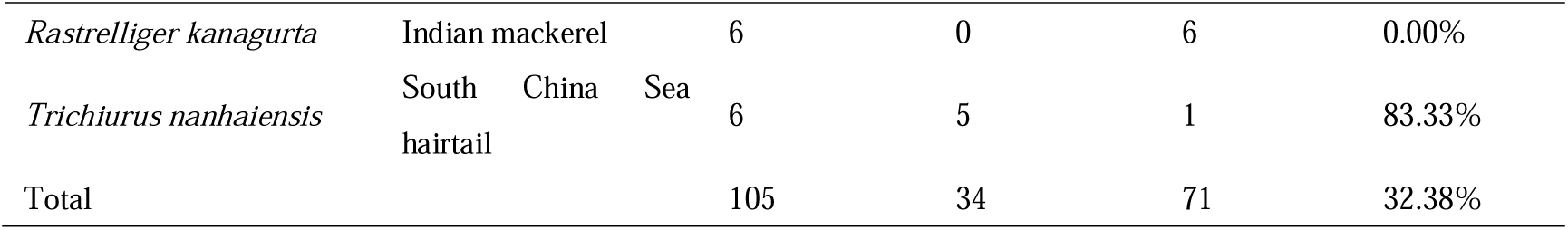
A summary of the distribution of *V. alginolyticus* isolates among the fish sampled for this study. Fish species were sampled from the Hong Kong waters of the South China Sea. The table lists the total number of individuals examined, positive and negative counts, and the corresponding positivity rate for each host species.

### 2.2 Isolation and culture of bacteria

In the laboratory, the dorsal fins of the collected fish were clipped, placed in a 15-mL tube with alkaline peptone broth (APB), and incubated at 37°C. After 24 h, the bacterial culture was streaked onto thiosulphate citrate bile salt sucrose (TCBS) agar with a loop and incubated at 37°C (first purification). After 24 h, three individual colonies, far from each other, were selected from each TCBS plate based on colony color and morphology. The three chosen colonies from each plate were transferred to a new TCBS plate by streaking and incubated at 37°C (second purification). After 24 h, single colonies from each isolate were placed in Eppendorf tubes with LB broth and incubated at 37°C for 24 h. Subsequently, 50% sterile glycerol, at the same volume as the bacterial culture, was added to each tube, and all samples were stored at −80°C for subsequent analysis (Fig. 1A).

**Figure 1.**
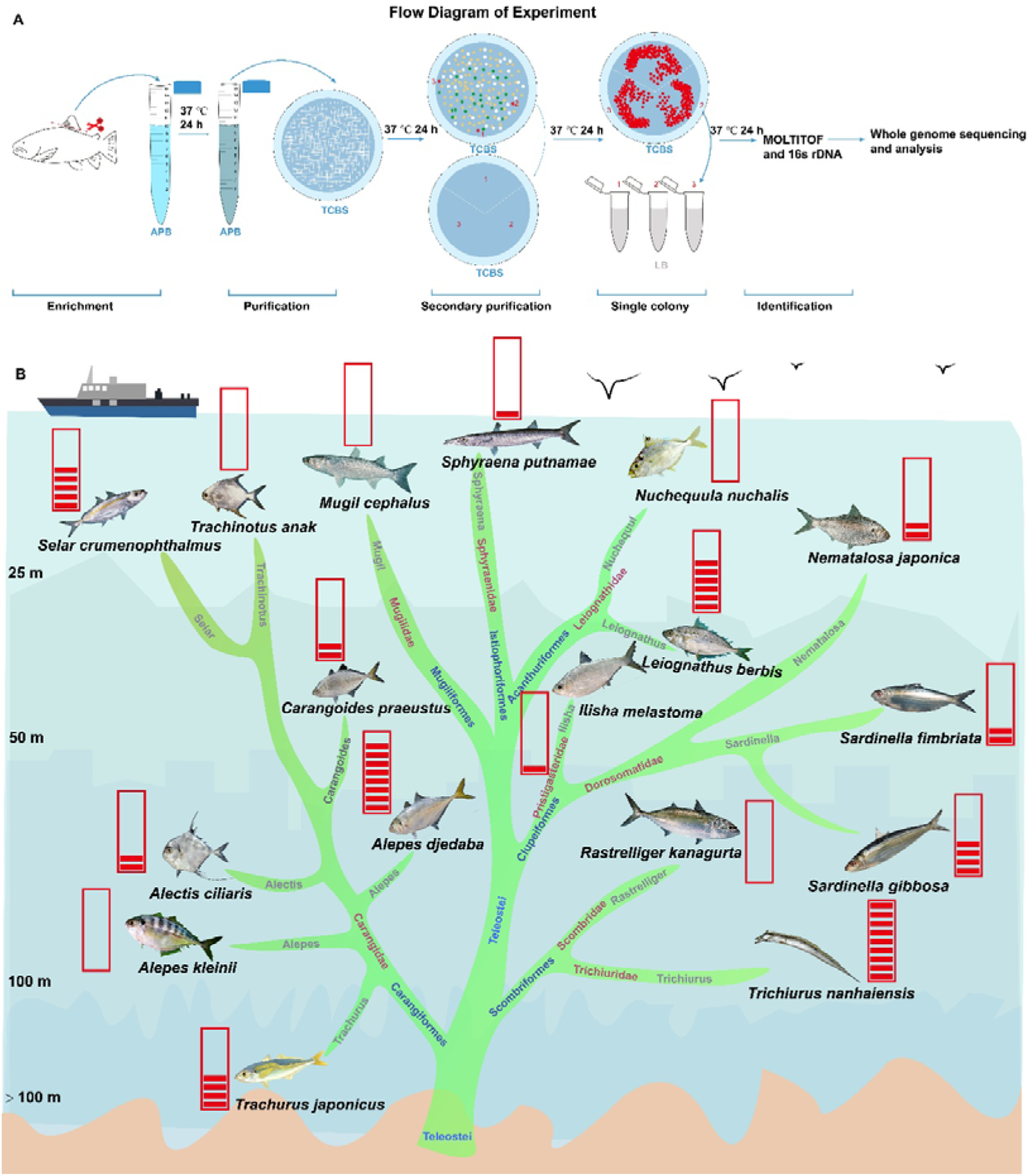
Isolation workflow and distribution of *V. alginolyticus* across wild marine fish species. (A) Schematic overview of bacterial isolation and analysis: enrichment culture, first purification, second purification, obtaining single colonies for storage, and identification. In the second purification, three colonies of *V. alginolyticus* were selected based on their color and morphology. (B) Abundance of *V. alginolyticus* isolates recovered from each fish species. Background shading indicates water depth strata: 0–25 m, 25–50 m, 50–100 m, and >100 m. Red bars represent the number of isolates per host species. Pictures of the fish were adapted from the FishBase website (https://www.fishbase.se/search.php).

### 2.3 Bacterial identification

The isolated bacteria, which were stored at −80 °C, were revived on LB agar plates. After incubation at 37°C for 24 h, single colonies were picked for identification by matrix-assisted laser desorption/ionization time-of-flight mass spectrometry (MALDI TOF MS) (Bruker Daltonics, Bremen, Germany). Species-level classification and spectral data analysis were performed using MALDI Biotyper v3.4 (Bruker Daltonics, Germany). Isolates achieving log (score) values ≥ 2.0 were identified at the species level according to manufacturer-recommended thresholds. Isolates with an identification result score value of less than 2 were further identified by 16S rRNA sequencing. The 16S rRNA was amplified with the bacterial universal primer pair 27F: AGAGTTTGATCMTGGCTCAG and 1387R: GGGCGGWGTGTACAAGGC (Riggio et al., 2014). The PCR mixture comprised 12.5 μL of 2× Rapid Taq Master Mix, 2 μL of bacterial template, 1 μL of each primer, and 8.5 μL of nuclease-free water. The PCR thermal cycling conditions were as follows: 4 min at 95 °C followed by 34 cycles of 30 s at 95 °C, 30 s at 55 °C, 1.5 min at 72 °C, and an extra extension step of 72 °C for 10 min. The amplified sequences were compared with the GenBank database (https://www.ncbi.nlm.nih.gov/) with BLAST analysis.

### 2.4 DNA extraction and sequencing

Genomic DNA was extracted from confirmed *Vibrio alginolyticus* strains using the TaKaRa MiniBEST Bacteria Genomic DNA Extraction Kit Ver. 3.0 (Takara, Japan), following the manufacturer’s standard protocols. The yield, purity, and integrity of purified genomic DNA were validated using a NanoDrop spectrophotometer (Thermo Fisher Scientific) and 1.5% agarose gel electrophoresis. For Oxford Nanopore long-read library construction, 1 μg of qualified genomic DNA was processed with the Ligation Sequencing gDNA-Native Barcoding Kit 24 V14 (SQK-NBD-114.24, ONT, Oxford Nanopore Technologies, UK). Whole-genome long-read sequencing was performed on a MinION (ONT) platform using R.10.4.1 Flow Cell (FLO-MIN114, ONT). The resulting pod5 files were base-called with Dorado v0.4.1 (https://github.com/nanoporetech/dorado) with the “dna_r10.4.1_e8.2_400bps_sup@v4.2.0” model.

### 2.5 *de novo* genome assembly and whole-genome analysis

Sequencing quality metrics of Oxford Nanopore long reads were summarized and visualized using NanoPlot v1.41.6. Prior to de novo genome assembly, simplex basecalled ONT reads were filtered with Filtlong, applying the parameters -min-mean-q 80 and -min_length 1000 to retain high-quality valid sequences. The obtained high-quality reads were subsequently assembled into draft genomes using Flye v2.9.4 (Zhang and Austin, 2005) with the default parameters. The assembled genomes were subsequently polished using long reads with Medaka v1.11.3 (https://github.com/nanoporetech/medaka). The completeness of polished assemblies was assessed via BUSCO (Benchmarking Universal Single-Copy Orthologs) v5.7.1 (Manni et al., 2021), with the vibrionales_odb10 database containing 1445 universal single-copy orthologs as the reference dataset. Genome-wide functional annotation was implemented using Prokka v1.14.6 (Seemann, 2014), in which protein-coding gene prediction was accomplished by Prodigal v2.6.3 (Hyatt et al., 2010). Furthermore, Gene Ontology (GO) functional classification and Kyoto Encyclopedia of Genes and Genomes (KEGG) pathway enrichment analyses were conducted using the ClusterProfiler package (Yu et al., 2012).

### 2.6 Pan-genome analysis

All whole-genome sequences were annotated by Prokka v1.14.6 (Seemann, 2014), and then the protein sequences were obtained. To reconstruct the pangenome of the 89 bacterial isolates, the predicted protein sequences derived from their genome assemblies were clustered into orthologous groups (orthogroups) using OrthoFinder v2.5.5 (Emms and Kelly, 2019) with default parameters. A U-shape distribution plot was constructed to illustrate gene frequencies, where orthogroups were classified into four categories based on their occurrence: the core genome (present in all 89 isolates), the soft-core genome (present in 82 to 88 isolates), the accessory genome (present in 2 to 81 isolates), and singletons (present in only 1 isolate). A pangenome rarefaction curve was generated using the R package micropan (Snipen and Liland, 2015) to evaluate the pangenome size, utilizing 50 random permutations to account for the effect of genome addition order. Additionally, a tile plot displaying the presence/absence matrix of the orthogroups was generated.

### 2.7 Phylogenetic tree

The dataset comprises 46 strains obtained from our study. Additionally, we also included all 44 strains with complete genome sequences from NCBI, with *V. harveyi* as the outgroup. The .faa files after annotation were analyzed with OrthoFinder v2.5.5 (Emms and Kelly, 2019) to identify 2836 single-copy ortholog genes. Multiple sequence alignment was performed on these genes using MUSCLE v5.1 (Edgar, 2022). Finally, uDance v1.6.4 (Balaban et al., 2024), utilizing RAxML v8 (Stamatakis, 2014) for maximum likelihood tree construction, was used to build a phylogenetic tree based on these aligned genes.

### 2.8 Antimicrobial resistance and virulence genes analysis

The polished genome assemblies were subjected to the identification of antibiotic resistance (AMR) genes and virulence genes. AMR genes were screened using the Resistance Gene Identifier (RGI) (Alcock et al., 2023) tool against the Comprehensive Antibiotic Resistance Database (CARD) with default arguments. For virulence gene profiling, VFanalyzer (Liu, B. et al., 2019) was used to query the Virulence Factor Database (VFDB) with default parameter settings to characterize potential virulence factors in the assembled genomes. The identified virulence and resistance genes were subsequently linked to their corresponding orthogroups to evaluate their distribution across the core and accessory genomes.

### 2.9 Identification of motifs and DNA methyltransferases

Modified methylation basecalling for 4mC, 5mC, and 6mA was performed using Dorado v0.7.2 (https://github.com/nanoporetech/dorado) with the dna_r10.4.1_e8.2_400bps_sup@v5.0.0_4mC_5mC@v1 and dna_r10.4.1_e8.2_400bps_sup@v5.0.0_6mA@v1 models. Previous studies have demonstrated that 6mA is the most prevalent modification in bacteria and impacts various biological processes (Lu et al., 2025; Ma et al., 2025). Therefore, for 6mA methylation identification, we used the analytical pipeline described by Zhang et al. (Zhang et al., 2026). Briefly, a whole-genome amplification (WGA) dataset with scarcely any DNA methylation modifications was generated as a negative control to eliminate spurious base modifications detected in whole-genome sequencing (WGS) datasets across all strains. High-confidence 6mA sites were defined as positions with methylation levels ≥ 25% higher in WGS compared to those in WGA (Zhang et al., 2026). The resulting high-confidence methylated sites were subsequently subjected to methylated motif discovery. Flanking sequences spanning 12 bp upstream and downstream of each 6mA site were extracted using BEDTools v2.29.1 (Quinlan and Hall, 2010). The STREME program within MEME Suite v5.5.7 (Bailey et al., 2015) was used to identify significantly enriched motifs (*P* < 0.05). In parallel, the motif-search module embedded in Modkit v0.3.1 (https://github.com/nanoporetech/modkit) was applied for motif screening with the –min-sites parameter set to 100. For 4mC and 5mC methylation identification, we utilized modkit to summarize the modified bases and find highly modified motif sequences that were enriched. The putative DNA methyltransferases (MTases) for all strains were identified using DNA_methylase_finder v1.0.1 (https://github.com/mtisza1/DNA_methylase_finder) (Tisza et al., 2023). To verify the reliability of the detected DNA methylation motifs in our isolates, correlation analyses between these conserved sequences and putative DNA MTases were further performed. The MTlinker function in DNA_methylase_finder v1.0.1 was employed to search against the proteomes to link the motifs and MTases of each strain. In addition, the candidate MTase proteins were also annotated via BLASTP v2.15.0 (Camacho et al., 2009) homology searching against curated MTase sequences retrieved from the REBASE database (Roberts et al., 2003). Homologous proteins with experimentally validated binding motifs were retained according to rigorous screening thresholds, including sequence identity > 30%, e-value < 10^−10^, and bit score > 100 (Zhang et al., 2026).

### 2.10 Discovery of mobile genetic elements and DNA methyltransferase distribution

The mobile genetic elements (MGEs), including insertion sequences (IS), prophages, plasmids, and integrative and conjugative elements (ICEs), were annotated. Insertion sequences were scanned by ISEScan v1.7.2.3 (Xie and Tang, 2017) with a hidden Markov model (HMM). Prophage sequences were identified by the PHASTER (Arndt et al., 2016) web server. The plasmid sequences were directly extracted from the genome sequences of the strains. The identification of ICEs was systematically resolved via an in silico de novo prediction workflow driven by the ICEfinder v1.0 (Liu, M. et al., 2019), integrating the core spatial clustering logic derived from IslandPath-DIMOB v1.0.0 (Bertelli and Brinkman, 2018). Then, the annotated MTase of each strain was matched against the validated BED files of insertion sequences, prophages, plasmids, and ICEs of each strain by BEDtools v2.29.1 (Quinlan and Hall, 2010) to aggregate molecular coordinates.

### 2.11 Motif enrichment in regulation and CDS regions of AMR and virulence genes

The start and end sites of the AMR and virulence genes were obtained, and the 200-bp regions located directly upstream of the start codons were defined as the putative regulatory zones (Chao et al., 2015; Martin et al., 2024). The exact regulatory coordinates were extracted via BEDtools v2.29.1 (Quinlan and Hall, 2010) with strand-specific parameters (-s) to ensure strict upstream alignment across both chromosomes. All identified motifs were mapped to the 200 bp upstream promoter regulatory regions and coding sequence (CDS) of individual virulence and AMR genes. Motif density (per kilobase) within these gene-specific regions was calculated, followed by Fisher’s exact test against the corresponding genome-wide background motif density to determine the motif enrichment.

### 2.12 Antimicrobial susceptibility test

The antibiotic susceptibility profiles of all isolated *V. alginolyticus* strains were determined via the standard Kirby–Bauer disc diffusion assay. Ten commonly used antibiotics with corresponding standard dosages were included in the susceptibility assessment, including Ampicillin (AMP, 10 μg/disc), Cefamezin (CZ, 30 μg/disc), Gentamycin (GM, 10 μg/disc), Erythromycin (ERY, 15 μg/disc), Norfloxacin (NOR, 10 μg/disc), Ciprofloxacin (CIP, 5 μg/disc), compound sulfamethoxazole SMZ/TMP (SXT, 23.75/1.25 μg/disc), Polymyxin B (PB, 300 μg/disc), Doxycycline (DOX, 30 μg/disc), and Tetracycline (TET, 30 μg/disc). Briefly, bacterial suspensions were evenly streaked onto Tryptic Soy Agar (TSA) plates, followed by the placement of antibiotic discs on the inoculated agar surface. After 24 h of incubation at 37 °C, the diameter of the bacterial growth inhibition zone was measured and recorded. Antibiotic resistance phenotypes of the isolates were classified into three grades (susceptible, intermediate, and resistant) in accordance with the criteria specified by the Clinical and Laboratory Standards Institute (CLSI, 2018). In addition, the multiple antibiotic resistance (MAR) index was calculated to evaluate the antibiotic contamination risk of these isolates. Isolates with a MAR index > 0.2 were defined as originating from high-risk sources of antibiotic contamination (Krumperman, 1983).

### 2.13 Statistical analysis

Statistical analyses were performed using IBM SPSS 20.0 (IBM Corporation, Armonk, NY, USA), and a *P*-value of < 0.05 was considered statistically significant. The Kruskal-Wallis test was applied to compare the distributions of AMR and virulence genes across different grouping categories.

## 3 Results

### 3.1 *V. alginolyticus* profiles isolated from 17 different species of wild fish in the South China Sea

A total of 105 wild marine fish representing 17 species, 8 families, and 6 orders were collected from Hong Kong waters of the South China Sea. These fish occupied diverse trophic levels and vertical habitat strata, ensuring representative sampling of the local marine fish community (Supplementary Table 1). The trophic levels varied among species, with the highest being Sawtooth barracuda (*Sphyraena putnamae*) (4.5 ± 0.0 se) and the lowest being Japanese gizzard shad (*Nematalosa japonica*) (2.4 ± 0.16 se). In addition, these species inhabited different vertical ocean layers, primarily the surface layer, with some species, such as the African pompano (*Alectis ciliaris*) and Japanese jack mackerel (*Trachurus japonicus*), and South China Sea hairtail (*Trichiurus nanhaiensis*), belonging to benthopelagic species.

*V. alginolyticus* was isolated and identified from 34 of the 105 fish, corresponding to an overall positivity rate of 32.38% (Table 1). Higher positivity rates were observed in *T. japonicus*, goldstripe sardinella (*Sardinella gibbos*), and *T. nanhaiensis* (66.67%, 66.67%, and 83.33%, respectively). No *V. alginolyticus* was recovered from five species, including Spotnape ponyfish (*Nuchequula nuchalis*), Razorbelly scad (*Alepes kleinii*), Oyster pompano (*Trachinotus anak*), Flathead grey mullet (*Mugil cephalus*), and Indian mackerel (*Rastrelliger kanagurta*) (Table 1). Further analysis of the distribution across fish species yielded 46 distinct *V. alginolyticus* isolates (Fig. 1B). The greatest number of isolates was obtained from *T. nanhaiensis* (nine isolates), followed by shrimp scad (*Alepes djedaba*), bigeye scad (*Selar Crumenophthalmus*), and goldstripe sardinella (*S. Gibbosa*) (Fig. 1B). As most of these fish species are commercially exploited, these findings highlight a potential exposure risk to *V. alginolyticus* for fishermen, seafood handlers, and consumers.

### 3.2 High-quality complete genome assembly of 46 *V. alginolyticus* isolates using nanopore sequencing

Long-read nanopore sequencing was employed to generate high-quality, complete circular genomes for all 46 *V. alginolyticus* isolates. Sequencing yielded a total of 15,185,230 sequences with an average depth of 200.78x and an N50 of 9,043 bp. Following data assembly, we successfully constructed one large and one small circular chromosome for all strains, with genome sizes ranging from 5.09 to 5.45 Mb (Fig. 3A). Most strains harbored one plasmid, and a subset contained none or up to two plasmids. The average number of coding sequences (CDS) per genome was 4,688.78 CDS (Supplementary Table 2).

To enable comparative analyses, all 43 publicly available complete genomes of *V. alginolyticus* were retrieved from NCBI (Supplementary Table 3), with a history ranging from 1971 to 2010. These isolates originated from various sources, including fish, shellfish, and environmental samples. Briefly, seven strains were isolated from diseased animals or were designated “pathogenic” in the original reports and were denoted as “pathogen” strains, while isolates from the environment or recovered from non-diseased organisms were designated “non-pathogen” strains. Those isolates covered a diverse geographical area including ten countries or regions, with the majority isolated from China (61/89) (Fig. 2A). The assembly results of those isolates were consistent with those of the strains obtained in this study, including the number of plasmids, genome size, and the number of CDS sequences. Notably, the genome sizes of several isolates from Germany were larger than those of the other strains (Fig. 2B). This study adds 46 high-quality complete genomes, expanding the global genomic resource for *V. alginolyticus* by approximately 51.7% and enabling robust pan-genomic, phylogenetic, and epigenomic comparisons.

**Figure 2.**
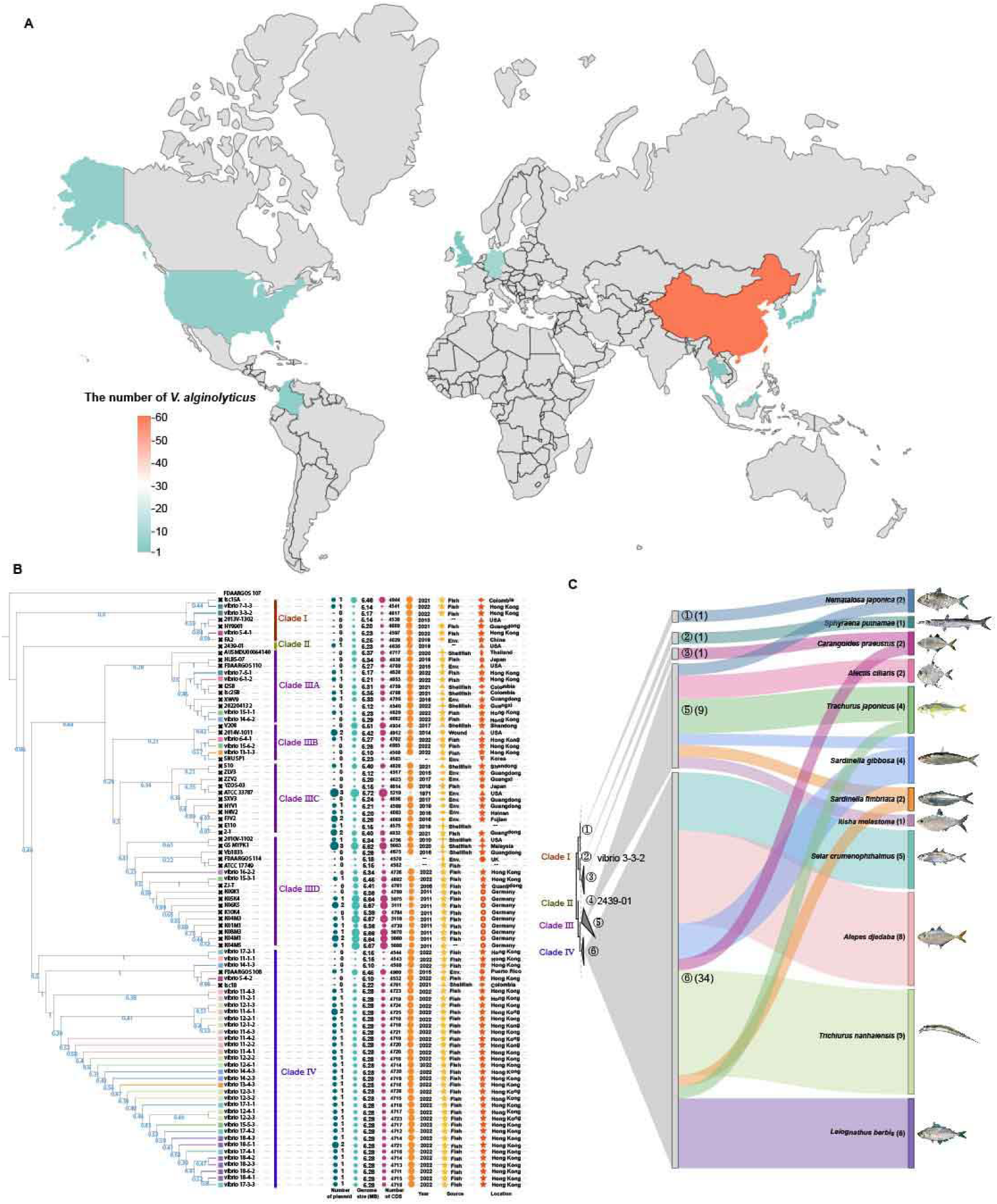
Phylogenetic tree and basic information on isolates of *V. alginolyticus*. (A) Global geographic distribution of 89 *V. alginolyticus* strains included in this study. The color shows the strain numbers per country/region. (B) The phylogenetic inferred from 2836 single-copy orthologue genes identified by OrthoFinder. The squares represent the strains obtained in this study, and crosses indicate NCBI reference strains. Clade assignment, plasmid count, genome size, CDS number, isolation year, source, and location are annotated. The bubble sizes are scaled for each panel, and the bubble sizes and different characters are not directly comparable across panels. Shellfish represent the strains isolated from shrimp and bivalves. (C) The phylogenetic tree was partitioned into six sections based on the distance matrix, showing the distribution of wild-fish isolates and their corresponding host species.

### 3.3 Phylogenetic clustering reveals a dominant host-shared clade (Clade) and broad genetic diversity of *V. alginolyticus* from wild fish

To further reveal the evolutionary relationships between these isolates, all *V. alginolyticus* strains, comprising 46 strains in this study and an additional 44 strains on NCBI with a *V. harveyi* as an outgroup, were used to construct a phylogenetic tree (Fig. 2B). Phylogenetic reconstruction using 2836 single-copy orthologous genes resolved 89 *Vibrio alginolyticus* genomes into four well-supported clades. Strikingly, 95.3% (41/43) of public reference strains were restricted to Clades I-III, while 73.9% (34/46) of isolates recovered from wild marine fish in this study fell exclusively within Clade IV, which greatly expanded the known genetic diversity of this clade. Geographical analysis revealed extensive intra-regional genetic heterogeneity. For example, strains from China, Colombia, and the USA were distributed across multiple clades, whereas German isolates formed a monophyletic group within Clade III. Isolates from the present study occupied all clades except Clade II, confirming their broad phylogenetic representation. This demonstrates that significant genetic diversity exists even within localized sampling regions (Fig. 2B).

Further analysis of host association patterns was conducted based on the distance matrix of the phylogenetic tree. Further partitioning of the tree by genetic distance showed that 73.9% (34/46) of wild-fish isolates, originating from 66.7% (8/12) of sampled fish species, clustered tightly within a single sub-lineage of Clade IV. Isolates from four demersal and reef-associated fish species (*Selar crumenophthalmus*, *Alepes djedaba*, *Trichiurus nanhaiensis*, and *Leiognathus berbis*) were entirely confined to this sub-lineage, indicating close evolutionary relatedness and frequent cross-species transmission. By contrast, isolates from most pelagic fish species (*C. praeustus*, *T. japonicus*, *S. fimbriata*, and *S. gibbosa*, except *N. japonica*) were scattered across multiple phylogenetic sections, suggesting that greater host mobility enhances environmental exposure and promotes the acquisition of genetically diverse strains. These results establish Clade IV as a distinct, ecologically specialized lineage strongly adapted to wild marine fish hosts, with phylogenetic structure shaped by host ecology and dispersal potential.

### 3.4 The pan-genome analysis reveals a closed gene repertoire and marked accessory genome depletion in Clade IV

Pan-genome analysis of 89 *V. alginolyticus* strains based on 421,234 annotated genes assigned 99.4% of sequences into 8,172 orthogroups (Supplementary Table 4). The pan-genome comprised 3,844 core gene families (47.03%) conserved across all strains, 269 soft-core gene families (3.29%) present in 82-88 strains, and 3,945 accessory gene families (48.27%) including strain-specific singletons (Fig. 3A). Rarefaction curves confirmed a closed pan-genome, indicating that the present dataset captures the near-complete gene repertoire of *V. alginolyticus* (Fig. 3B). Further integrated analysis of the phylogenetic tree and pan-genome revealed distinctive characteristics in clade IV. The pan-genome in this clade exhibited significant divergence from other clades. Core genomes in clades I-III were enriched in multi-copy gene clusters, whereas clade IV exhibited severe depletion of these features. Meanwhile, a large number of gene families in Clade IV within its accessory genome were not identified (Fig. 3C). Comparative analysis showed that Clades I and IV shared 4,801 gene families, with Clade I harboring 763 unique families versus 685 in Clade IV. Clades III and IV shared 5,349 gene families, consistent with a closer evolutionary relationship; however, Clade III encoded 2,588 unique gene families while Clade IV possessed only 137, representing a nearly 19-fold reduction (Fig. 4D). This also suggests that the gene clusters in the accessory genome of Clade IV seemed to be pronouncedly depleted.

**Figure 3.**
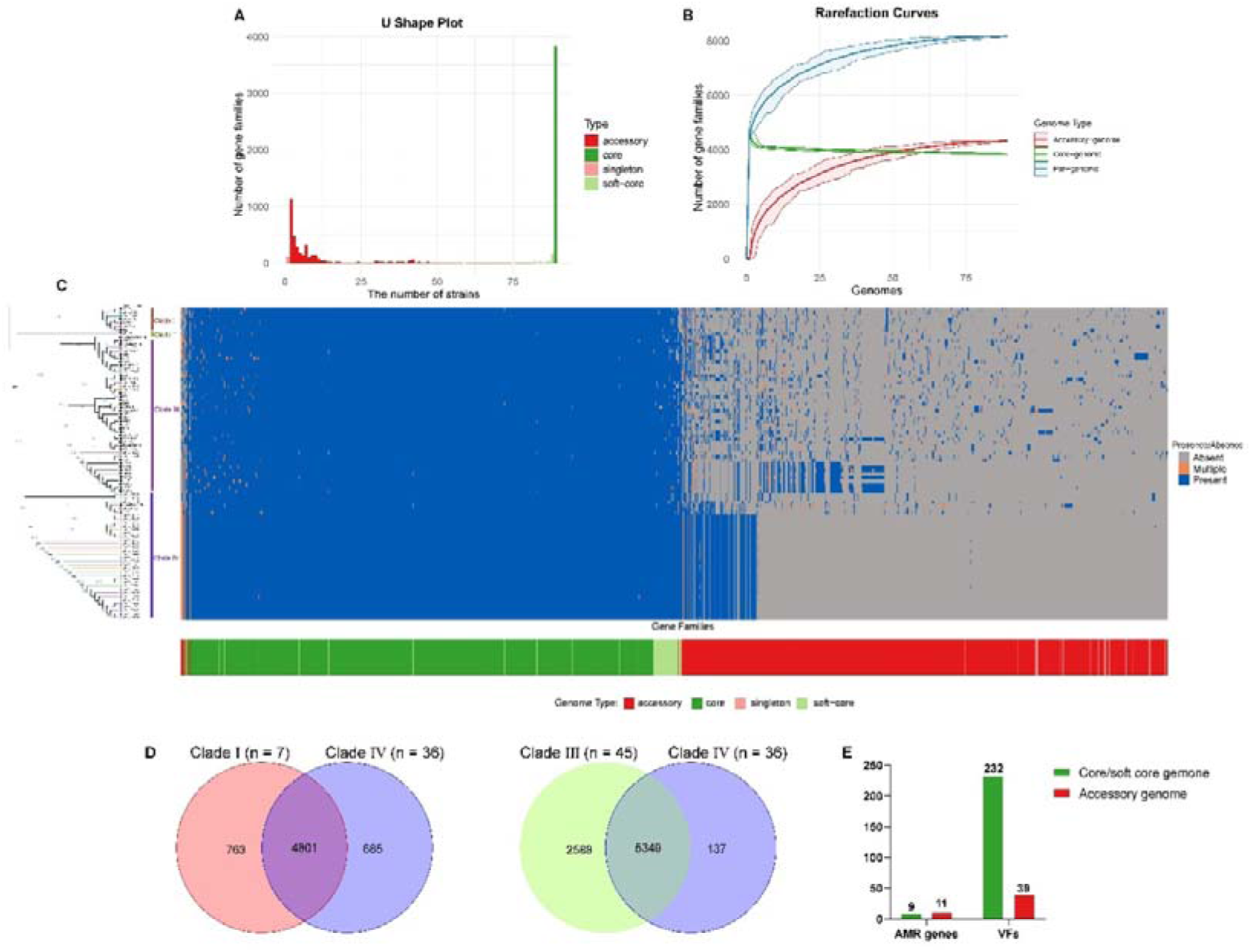
Pan-genome analysis of *V. alginolyticus* isolates. (A) Proportions of core, soft-core, accessory, and singleton gene families across 89 strains. (B) Rarefaction curve confirming a closed pan-genome. (C) Distribution and copy-number status of gene families within core, soft-core, and accessory compartments. (D) The Venn diagrams comparing gene families of Clades I and Clade IV, Clade III and Clade IV, respectively. (E) The distribution of antimicrobial resistance and virulence factor genes in the core and accessory genomes.

**Figure 4.**
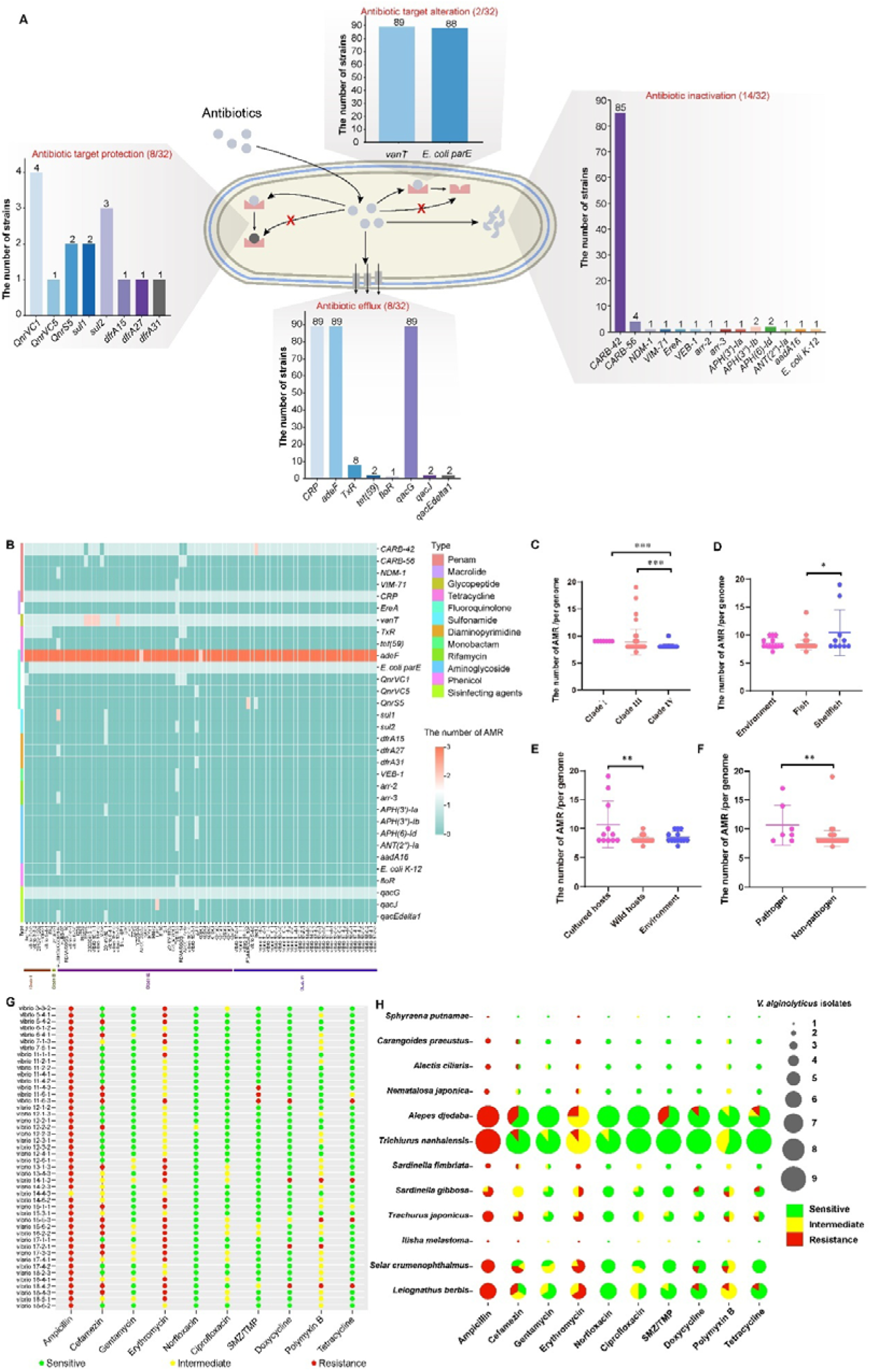
Antimicrobial resistance gene profiles and phenotypic resistance in *V. alginolyticus*. (A) The distribution of AMR genes in different resistance mechanisms. (B) The distribution of AMR genes across all strains. Thirty-two AMR genes exhibited resistance to 12 different antibiotics. (C-F) Comparisons of AMR gene counts by phylogenetic clade, isolation source, host type (wild vs aquaculture), and pathogenic status. Boxes show medians and interquartile ranges; whiskers denote minima and maxima. (G) Phenotypic resistance profiles of 46 wild-fish isolates against 10 antibiotics. (H) Antimicrobial susceptibility patterns of isolates stratified by host fish species. Circle size indicates isolate number; color indicates resistance phenotype.

To elucidate the functions of accessory genomes, we conducted Gene Ontology (GO) and Kyoto Encyclopedia of Genes and Genomes (KEGG) enrichment analyses. Functional enrichment analysis revealed that core genome genes were predominantly involved in fundamental metabolism, biosynthesis, and cellular processes such as oxidative phosphorylation and biofilm formation (Fig. S1A, S1B). Soft-core genes were enriched in signal transduction, xenobiotic metabolism, and drug metabolism pathways (Fig. S1C, S1D). By contrast, accessory genome genes were significantly enriched in negative regulation of metabolic pathways, defense responses, and biosynthesis of O-antigen sugars and polyketides, supporting key roles in environmental adaptation (Fig. S1E, S1F). Together, these findings demonstrate that Clade IV carries a streamlined accessory genome and represents a genetically distinct lineage shaped by unique evolutionary pressures.

### 3.5 Wild fish-derived *V. alginolyticus* strains in Clade IV exhibit a significantly reduced antimicrobial resistance gene repertoire

Genomic screening of antimicrobial resistance (AMR) determinants was performed on these 89 *V. alginolyticus* genomes. This screening identified 20 AMR-related gene families, comprising a total of 763 individual AMR genes that corresponded to 32 distinct AMR gene types. All identified AMR determinants were further categorized into four canonical functional mechanisms of antibiotic resistance: target protection, target alteration, antibiotic inactivation, and antibiotic efflux (Fig.3E, Fig. 4A, and Supplementary Table 5). The resistance genes in the core genome were primarily associated with resistance to antibiotics such as β-lactams, tetracyclines, glycopeptides, and fluoroquinolones, indicating that *V. alginolyticus* may have a natural resistance to these specific antibiotics. Conversely, the resistance genes in the accessory genome exhibited resistance to these major antibiotic classes and several additional antibiotics, including sulfonamides, rifampicin, and aminoglycosides (Supplementary Table 5).

Of these, six core AMR genes were universally conserved, including *CARB-42/CARB-56* (inactivation), *CRP, adeF, qacG* (efflux), *vanT*, and *E. coli parE* (target alteration), with *adeF* present in multiple copies (Fig. 4A and 4B), indicating that inactivation, efflux, and target alteration represent the primary intrinsic resistance strategies in this species. Notably, nearly all AMR genes were chromosomally encoded, and only a single strain (2014V-1011) carried resistance determinants on plasmids (Fig. S2). In addition, it was found that some strains exhibited a broader spectrum of AMR genes, such as AUSMDU00064140, Vb1833, and ZJ-T, which were isolated from cultured animals in Thailand and China and were clustered in Clade III (Fig. 4B). Subsequently, comparative analysis revealed that Clade IV isolates harbored a significantly lower abundance of AMR genes compared with strains in Clades I and III (Fig. 4C). A strong host-associated pattern was also observed: isolates recovered from wild fish carried fewer AMR genes than those from shellfish, aquaculture hosts, or designated pathogenic strains (Fig. 4D-4F). However, it showed no significant relationship between AMR genes and location, isolation time, and fish species (Fig. S3A-S3C).

Consistent with genomic predictions, phenotypic antimicrobial susceptibility testing of the 46 wild-fish isolates showed high resistance to ampicillin (97.82%) and erythromycin (52.17%), full sensitivity to gentamicin, norfloxacin, and ciprofloxacin, and low resistance rates (<10%) to cefamezin, SMZ/TMP, doxycycline, tetracycline, and polymyxin B (Table 2 and Fig. 4G). The multiple antibiotic resistance (MAR) index ranged from 0 to 0.6, with 32.61% (15/46) of isolates classified as multidrug-resistant (MAR > 0.2) (Supplementary Table 6). Multidrug-resistant strains were widely distributed across host species, with the highest resistance phenotypes (MAR = 0.6) detected in isolates from *A. djedaba* and *L. berbis* (Fig. 4H and Fig. S4). Together, these results indicate that intensive antibiotic selection in aquaculture drives elevated AMR, whereas wild fish–associated *V. alginolyticus* (particularly those in Clade IV) retain a low-resistance phenotype consistent with limited anthropogenic antibiotic exposure in wild fish.

**Table 2.** Distribution of antimicrobial susceptibility of *V. alginolyticus* isolated from different fish species in this study

| Antibiotics | Category | Sensitive | Intermediate | Resistant | Resistance | Predicted genes |
| --- | --- | --- | --- | --- | --- | --- |
| Ampicillin (10 µg) | Penam | 0 | 1 | 45 | 97.82% | 100% |
| Cefamezin (30 µg) | Cephalosporins | 21 | 10 | 15 | 32.61% | 0% |
| Gentamycin (10 µg) | Aminoglycoside | 35 | 11 | 0 | 0.00% | 0% |
| Erythromycin (15 µg) | Macrolide | 0 | 22 | 24 | 52.17% | 100% |
| Norfloxacin (10 µg) | Fluoroquinolone | 45 | 1 | 0 | 0.00% | 100% |
| Ciprofloxacin (5 µg) | Fluoroquinolone | 34 | 12 | 0 | 0.00% | 100% |
| Sulfamethoxazole/Trimethoprim SMZ/TMP (23.75/1.25 µg) | Sulfonamides | 40 | 3 | 3 | 6.52% | 0% |
| Doxycycline (30 µg) | Tetracycline | 41 | 1 | 4 | 8.70% | 6.52% |
| Tetracycline (30 µg) | Tetracycline | 40 | 2 | 4 | 8.70% | 6.52% |
| Polymyxin B (300 µg) | Polypeptide | 18 | 24 | 4 | 8.70% | 0% |

### 3.6 Virulence gene profiling reveals a depauperate virulence repertoire in wild fish-associated Clade IV

Virulence factor (VF) screening across the pan-genome identified 271 VF-associated gene families, of which 232 (85.61%) belonged to the core or soft-core genome, while 39 (14.39%) were located in the accessory genome (Supplementary Table 7 and Fig. S5). This indicates that most virulence determinants are stably inherited and essential for the lifestyle of *V. alginolyticus*. Isolates encoded a conserved suite of canonical Vibrio virulence modules, including genes associated with adhesion, secretion systems, toxin production, and quorum sensing. Notably, all strains harbored a complete T3SS1 gene cluster identical to that of *V. parahaemolyticus*, suggesting shared pathogenic mechanisms (Fig. 5A). Virulence loci were unevenly distributed between the two chromosomes: genes for adhesion, chemotaxis, motility, and secretion systems were located exclusively on chromosome 1 (the bigger one), whereas functions related to antiphagocytosis and iron uptake were concentrated on chromosome 2 (the smaller one). Plasmids carried almost no virulence genes, except *pilW* in strain ATCC_33787 (Fig. 5A). All isolates tested positive for the thermolabile hemolysin gene *tlh*, but lacked the high-risk human pathogenic determinants *tdh* and *ctxB*, indicating a reduced zoonotic potential relative to other pathogenic Vibrio species. Some strains still possessed virulence factors such as *ctrD*, *cysE*, and *wbtI*, which are associated with immune evasion, along with endotoxin genes like *bplF*, *bplG*, and *kpsF* (Fig. 5A). These genes were also identified in the isolates examined in this study, indicating that the universal presence of these virulence modules warrants caution regarding *V. alginolyticus* exposure during aquaculture and recreational marine activities.

**Figure 5.**
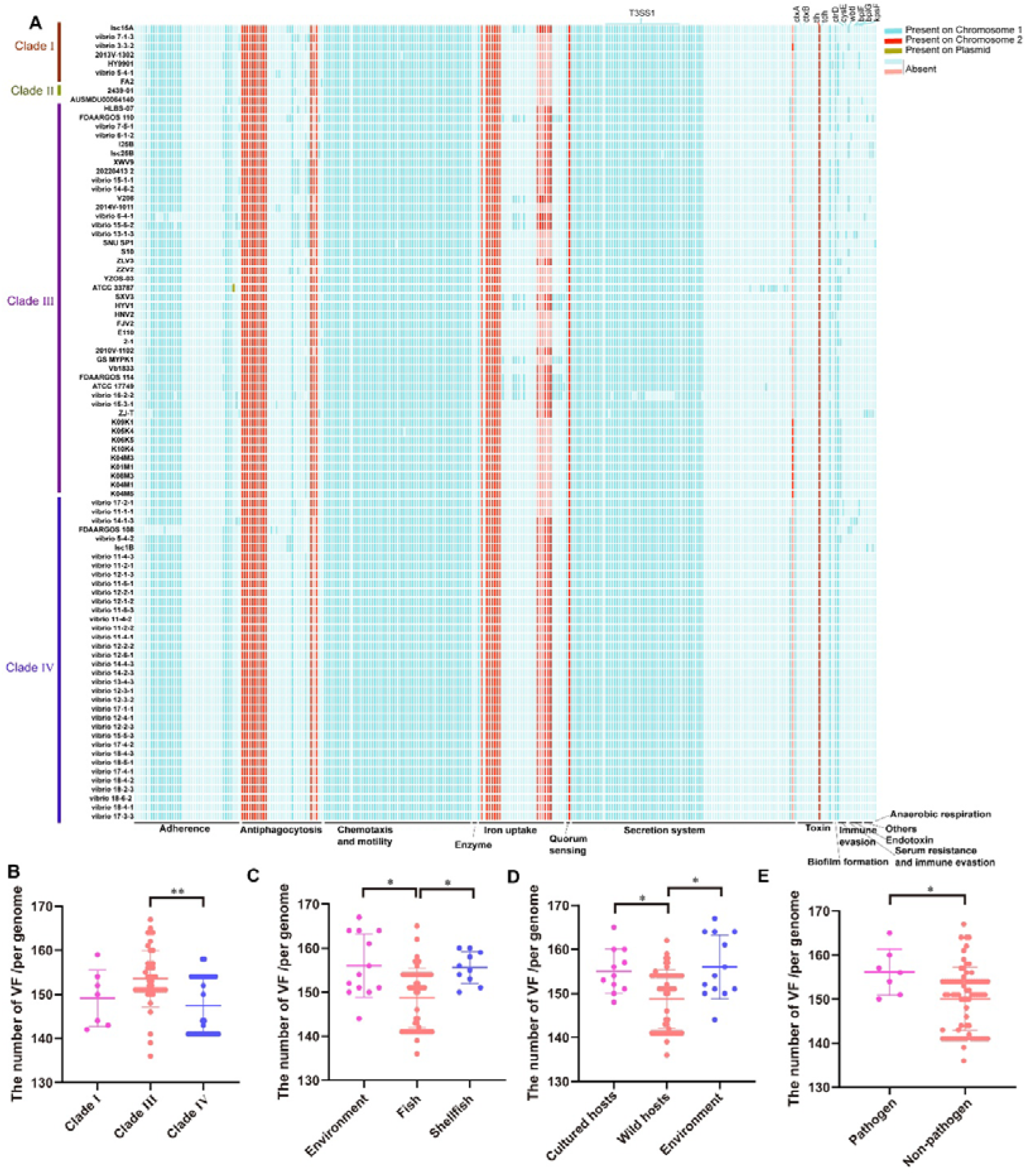
Virulence gene composition and distribution in *V. alginolyticus*. (A) The distribution of different virulence genes in the genomes of *V. alginolyticus*. Long bars of various colors indicated the presence of virulence genes in a strain and specified their chromosomal locations. The virulence gene classes were listed at the bottom. (B-E) Comparisons of virulence gene counts by phylogenetic clade, isolation source, host type (wild vs aquaculture), and pathogenic status.

Parallel to the AMR profile, Clade IV isolates carried a significantly lower abundance of virulence genes compared with Clade III (Fig. 5B). Isolates from wild fish also displayed a markedly reduced virulence gene repertoire relative to those from shellfish, aquaculture environments, and pathogenic strains (Fig. 5C-5E). These patterns support an evolutionary trade-off: *V. alginolyticus* in wild fish maintains a low-virulence profile to sustain commensal persistence, whereas aquaculture-associated lineages accumulate enhanced virulence traits under strong host- and environment-driven selection. In addition, South American isolates carried a significantly higher virulence gene burden than European isolates (Fig. S6A). However, no significant association was observed between virulence gene counts and isolation year or host fish species, although minor variation was noted among individual isolates (Fig. S4B and S4C).

### 3.7 Plasmidome analysis reveals plasmids contribute to environmental adaptation rather than virulence or antibiotic resistance

It has been shown that in bacteria, plasmids carry resistance and virulence genes that confer antibiotic resistance and virulence to strains. However, this study found that some *V. alginolyticus* strains did not contain plasmids, and in strains that did contain plasmids, the plasmids carried very little virulence (except for the strain ATCC_33787) and resistance genes (except for the strain 2014V-1011) (Fig. S2 and Fig. 5A). This suggests that plasmids may be involved in other physiological functions in *V. alginolyticus*. Consequently, we conducted a further analysis of the plasmidome of *V. alginolyticus*. A total of 65 complete circular plasmids were identified across 56 *Vibrio alginolyticus* strains, with sizes ranging from 9,064 bp to 300,425 bp and an average of 141.8 coding sequences per plasmid (Supplementary Table 8). Functional enrichment analysis showed that plasmid-encoded genes were significantly associated with stress responses, DNA replication, recombination, repair, and core metabolic processes (Fig. S7A). KEGG enrichment further highlighted functions related to bacterial defense mechanisms, DNA mismatch repair, and homologous recombination (Fig. S7B). These results demonstrate that *V. alginolyticus* plasmids play a minor role in pathogenicity and drug resistance, instead functioning as accessory modules that enhance environmental adaptability, genome plasticity, and survival under fluctuating marine conditions.

### 3.8 Mobile genetic elements mediate the diversification and horizontal dissemination of accessory genome-associated methyltransferases

DNA methyltransferases are believed to play a crucial role in regulating various physiological functions in bacteria (Oliveira and Fang, 2021). Therefore, we further analyzed the methyltransferases. We identified 355 DNA methyltransferase genes belonging to 29 gene families across 88 *V. alginolyticus* strains, covering Type I, Type II, Type IIG, Type III, and unclassified methyltransferases (Fig. 6A and Supplementary Table 9). Each genome contained 1-12 methyltransferase genes, with 71.90% (255/355) located in the accessory genome, indicating high epigenetic diversification across lineages. Furthermore, we characterized the distribution of these MTases across mobile genetic elements (MGEs), including IS, plasmids, ICEs, and prophages. In total, 31.6% (106/335) MTases were found within MGEs, all of which were located in the accessory genome (Supplementary Tables 9 and 10). Plasmids were the dominant vector for horizontal transfer of MTases, carrying 68.9% of MTases (73/106 of all MGE-derived MTases), followed by prophages (47 MTases, 44.3%), while ICEs only carried 15.1% (16/106) MTases. Notably, 21.7% (23/106) MTase genes resided in overlapping regions of nested MGEs and were annotated to two distinct element types (plasmid/prophage, plasmid/ICE, or ICE/prophage). These findings indicate that horizontal gene transfer serves as a crucial route for Vibrio to acquire DNA methyltransferase genes, with plasmids acting as the dominant vehicle for MTase dissemination among strains. The widespread overlapping localization also suggests frequent recombination and nested integration between distinct mobile elements, enabling the shuttling of MTase modules across different MGE backbones. In addition, the acquisition of MTases facilitates MGE evasion from the host’s endogenous restriction system and promotes stable maintenance inside host genomes.

**Figure 6.**
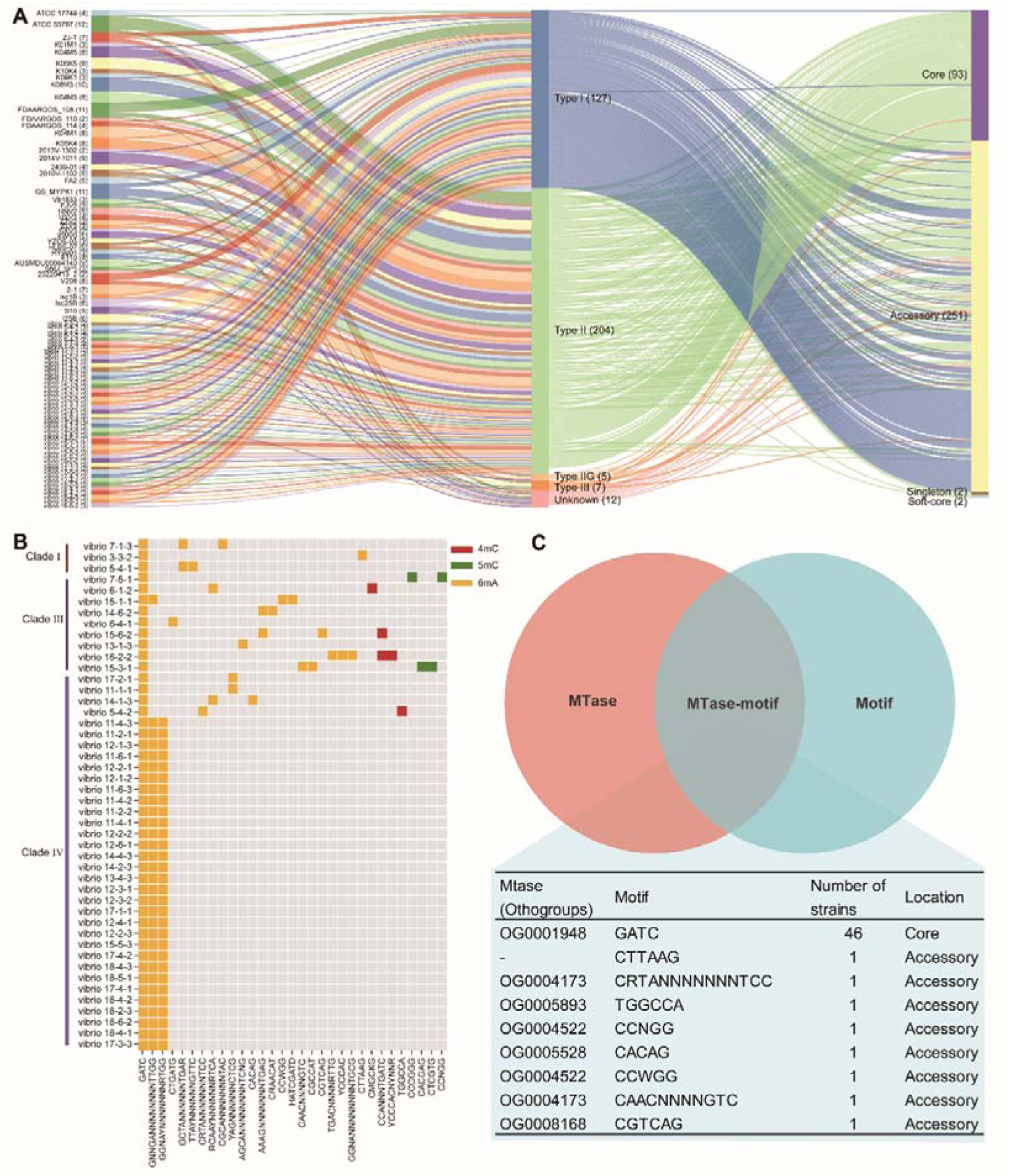
DNA methyltransferases and methylation motifs. (A) Abundance, type, and pan-genomic localization of DNA methyltransferases (MTase). Total counts and genomic distribution (core, soft-core, accessory, singleton) of Type I, II, IIG, III, and unclassified DNA methyltransferases across all *V. alginolyticus* strains. Numbers in parentheses indicate total methyltransferase genes per strain. (B) Strain-specific DNA methylation motifs in wild-fish *V. alginolyticus*. Distribution of 6mA-, 4mC-, and 5mC-associated methylation motifs across 46 isolates, grouped by phylogenetic clade. Most motifs are strain-specific, and Clade IV uniquely shares a conserved subset. (C) The Venn diagram of DNA methyltransferases and motifs.

The intersection represents the matched pairs of DNA methyltransferases and their corresponding methylation motifs. The table lists all MTase-motif pairs identified via the MTlinker pipeline. Only the GATC motif was matched to its cognate DNA methyltransferase in all strains.

### 3.9 Conserved and strain-specific methylation motifs exhibit distinct enrichment profiles in virulence and antimicrobial resistance genes

Using nanopore long-read sequencing, we systematically detected 6mA, 4mC, and 5mC DNA modifications and identified 140 methylation motifs grouped into 31 distinct families in the 46 wild-fish isolates (Fig. 6B and Supplementary Table 11). The 6mA modification dominated (93.57%), with the canonical GATC motif present in all strains, consistent with previous studies on prokaryotes ^(Seong^ ^et^ ^al., 2021)^, while 4mC and 5mC methylated motifs constituted only 3.57% and 2.86%, respectively. Most remaining motifs were strain-specific, including unique 6mA, 4mC, and 5mC motifs detected in only one isolate, such as “CTGATG”, “TTAYNNNNNNGTTC,” and “CACAG” in 6mA modification, “CMGCKG” and “TGGCCA” in 4mC modification, and “CCCGGG” and “CACGAG” in 5mC modification. We identified 17 distinct methylation motif combinations, which were largely strain-specific and showed no consistent correlation with the phylogenetic tree (Fig. S8). A conserved combination of three motifs (GATC, GNNGANNNNNNNNTTGG, and GGNAYNNNNNNNRTGG) was exclusively shared by Clade IV strains, providing an epigenetic signature that supports the evolutionary coherence of this wild-fish-adapted clade.

We further employed the MTlinker function in DNA_methylase_finder to link identified DNA methyltransferases with their corresponding methylation motifs across isolates. Only a limited number of motifs could be matched to cognate methyltransferases, such as “CTTAAG”, “TGGCCA”, and “CCNGG”, which were also identified in one strain. Notably, the universal core motif “GATC”, which was detected in all strains, was successfully annotated with its corresponding methyltransferase in all isolates (Fig. 6C and Supplementary Table 12). The gene encoding this GATC-specific methyltransferase is located within the core genome, highlighting the essential role of this conserved methyltransferase–motif pair in the survival and fundamental physiological homeostasis of *V. alginolyticus*.

To characterize the distribution of DNA motifs associated with potential regulatory and coding functions, we performed motif enrichment analysis on the coding sequences (CDS) and regulatory regions (200 bp upstream of CDS) of AMR and virulence genes (Supplementary Tables 13-16). For AMR genes, CDS and regulatory regions presented distinct motif enrichment patterns. The *qacG* CDS showed consistent significant motif (GGNAYNNNNNNNRTGG) enrichment across most strains (14/46) (*P* < 0.05), while other AMR genes only exhibited sporadic, strain-specific enrichment (Fig. S9 and Supplementary Table 13). In regulatory regions, *vanT* was the core enriched locus, with a single motif (GNNGANNNNNNNNTTGG) significantly enriched across 16/46 strains (*P* < 0.05) (Fig. S10 and Supplementary Table 14), and no shared motifs with CDS regions. These results indicate that AMR genes have distinct sequence methylation preferences in coding and regulatory domains, with no universal sequence methylation features across different AMR loci. For virulence genes, the GATC motif was significantly enriched both in CDS and regulatory regions covering multiple functional gene families, including flagellar structural genes (*flg/flh/fli*), toxins, secretion systems, and polysaccharide biosynthesis (*cps*) (*P* < 0.05) (Fig. S11 and S12). Many strains showed significant enrichment signals across multiple virulence genes, reflecting strong and consistent motif accumulation. The enrichment pattern varied considerably between strains, with some strains displaying broad GATC enrichment across numerous virulence loci, while others showed only limited signals, suggesting that strain-specific genomic variations may influence motif distribution. Non GATC motifs displayed extremely restricted, strain specific, and sparse enrichment in either regulatory or coding regions of virulence genes (Supplementary Tables 15 and 16). Additionally, prominent strain heterogeneity was observed for motif enrichment across both AMR and virulence genes, which reflected epigenetic variation among individual *Vibrio* isolates. These findings demonstrate that the essential GATC motif potentially mediates epigenetic modification of virulence genes in *V. alginolyticus*, whereas it exerts no such effect on AMR genes.

## 4 Discussion

*V. alginolyticus* is a globally distributed opportunistic pathogen endemic to estuarine and marine environments, and represents the second most common *Vibrio* species associated with human clinical infections (CDC, 2012). It also causes severe disease and mass mortality in a wide range of cultured aquatic animals, leading to substantial economic losses in mariculture worldwide (Cao et al., 2018; Deng et al., 2020; Rameshkumar et al., 2017; Yang et al., 2021). Despite its clinical, veterinary, and ecological importance, large-scale genomic and epigenomic investigations of *V. alginolyticus* from wild fish hosts remain scarce. Most previous studies have focused on diseased aquaculture specimens, leaving the prevalence, population structure, virulence potential, and antimicrobial resistance profiles of *V. alginolyticus* in wild marine ecosystems poorly understood. In this study, we performed systematic sampling of 105 wild marine fish representing 17 species from Hong Kong waters of the South China Sea, yielding 46 *V. alginolyticus* isolates with a prevalence of 32.38%. Although no *V. alginolyticus* was isolated from some species, more isolates were detected in some important commercial fish species, such as *T. japonicus* (66.67%), *S. gibbosa* (66.67%), and *T. nanhaiensis* (83.33%), indicating potential health risks for fishermen, seafood handlers, and consumers. Using high-coverage nanopore long-read sequencing, we generated complete circular genomes and DNA methylome profiles for these strains, and integrated them with all publicly available complete genomes of *V. alginolyticus* to establish the most comprehensive pan-genomic, resistomic, virulomic, and epigenomic framework for this species to date.

The construction of phylogenetic trees offers a robust framework for interpreting evolutionary relationships among *V. alginolyticus* strains. It is well-established that strains clustering within the same clade share closer genetic relatedness and likely descend from a recent common ancestor (Huang et al., 2024). Phylogenomic analysis resolved 89 *V. alginolyticus* strains into four well-differentiated clades. Strikingly, 95.3% of public reference strains were confined to Clades I-III, whereas 73.9% of isolates from wild fish in this study clustered within Clade IV, indicating that this clade represents a largely uncharacterized wild-fish-adapted lineage. Geographical distribution showed no strict phylogeographic structure, consistent with previous reports that *V. alginolyticus* exhibits high genetic diversity within local populations and undergoes frequent cross-regional transmission (Huang et al., 2024). Further partitioning revealed that most wild-fish isolates clustered tightly within a single sub-lineage of Clade IV, with strains recovered from eight different fish species, indicating broad host sharing and potential cross-species transmission. Isolates from demersal and reef-associated fish were phylogenetically cohesive, whereas those from pelagic fish displayed greater diversity, likely reflecting increased environmental exposure and dispersal driven by host mobility. These results reveal that host ecology and behavior shape the population structure of *V. alginolyticus* more strongly than geographic isolation.

The pan-genome provides a comprehensive representation of a species’ genetic diversity, overcoming the limitations inherent in using a single reference genome (Tettelin et al., 2005). By integrating multiple individual genomes, it elucidates the complete gene repertoire of a species and its associated variations, thereby offering a crucial foundation for research in agricultural breeding, disease mechanisms, species evolution, and adaptive dynamics (He et al., 2025; Muzzi et al., 2007; Yin et al., 2024). Pan-genome analysis of 89 strains yielded a closed pan-genome composed of 3,844 core gene families (47.03%), 269 soft-core families (3.29%), and 3,945 accessory gene families (48.27%). A closed pan-genome indicates that our dataset captures the near-complete gene repertoire of *V. alginolyticus*, which provides a robust foundation for evolutionary and functional studies. Notably, Clade IV exhibited substantial depletion of accessory genome content and multicopy gene clusters compared with Clades I-III, suggesting genome streamlining and niche specialization in wild fish hosts. Functional enrichment showed that core genes support essential cellular processes, while accessory genes contribute to environmental adaptation, defense, and surface polysaccharide biosynthesis. The reduced accessory genome in Clade IV likely reflects a commensal lifestyle in wild fish, where minimal metabolic burden and stable host colonization are favored over extensive genomic plasticity. This pattern contrasts with many marine pathogens, in which open pan-genomes facilitate rapid adaptation to fluctuating aquaculture environments.

Plasmids are extrachromosomal DNA molecules capable of autonomous replication and stable coexistence with host chromosomes. They have emerged as key drivers of horizontal gene transfer (HGT), playing a fundamental role in prokaryotic evolution. Recent studies further reveal that plasmid-borne genes evolve through distinct mechanisms compared to chromosomal genes, indicating that plasmids function not merely as passive vectors for gene exchange, but as dynamic evolutionary entities in their own right (Rodríguez-Beltrán et al., 2021). Understanding these molecular dynamics is essential for predicting and addressing the spread of critical plasmid-encoded traits, such as antimicrobial resistance and bacterial virulence (Lassalle et al., 2023; Wang et al., 2025; Wang et al., 2022). For example, studies confirmed that *V. harveyi* harbored up to five distinct plasmids, all of which correlate strongly with enhanced strain virulence. Strains carrying these plasmids demonstrate significantly higher host mortality compared to plasmid-free strains (Wang et al., 2025). These results underscore the pivotal role of plasmids in disseminating antibiotic resistance and pathogenicity throughout the *Vibrio* genus. In this study, plasmidome analysis identified 65 plasmids across 56 strains, but these elements carried almost no virulence or antimicrobial resistance determinants. Instead, plasmid-encoded genes were enriched in stress response, DNA replication, recombination, and repair, which indicated that plasmids contribute to environmental adaptability and genome stability rather than pathogenicity and resistance. In most *Vibrio* species, including *V. cholerae* and *V. parahaemolyticus*, plasmids are major vectors for horizontal transfer of resistance and virulence genes (Lassalle et al., 2023; Wang et al., 2022). Our findings reveal a distinct evolutionary strategy in *V. alginolyticus*, in which key fitness determinants are stably maintained on the chromosome, while plasmids serve as accessory modules for survival in dynamic marine habitats. This architecture reduces the risk of horizontal transmission of hypervirulent or multidrug-resistant lineages but ensures robust persistence in coastal ecosystems.

Recently, bacterial antibiotic resistance has become a significant public health concern. Since the discovery of penicillin by Alexander Fleming in 1928, more and more antibiotics have been given wide application in various areas, such as medicine and agriculture (Durand et al., 2019; Zhang et al., 2021). Resistance mechanisms usually include mutation, target site protection, antibiotic efflux, and antibiotic inactivation (Van Goethem et al., 2018). In *Vibrio* species, virtually all of these strategies were employed (Nathamuni et al., 2019), which reflects a remarkable diversity of resistance mechanisms that underscored their adaptive versatility and posed a significant challenge for clinical and environmental management. In addition, among clinical *Vibrio* isolates, many have been reported to be resistant to common antibiotics, which would hamper effective treatment against these pathogens (Elmahdi et al., 2016). Multiple drug resistance has also been reported in environmental *Vibrio* isolates (Igbinosa, 2016). Similarly, some *V. alginolyticus* isolates from coastal aquaculture areas have shown multidrug resistance (Yu et al., 2022). In the present study, antimicrobial resistance profiling identified 32 AMR gene types belonging to four mechanisms, with efflux pumps and antibiotic inactivation as the dominant strategies. Six core resistance genes were conserved across all strains, conferring intrinsic resistance to β-lactams, macrolides, and fluoroquinolones. Consistent with genomic predictions, phenotypic assays showed that 97.82% of wild-fish isolates were resistant to ampicillin, 52.17% to erythromycin, and fully susceptible to gentamicin, norfloxacin, and ciprofloxacin. Ampicillin belongs to the β-Lactam antibiotics, the most common and widely used group (Mitchell et al., 2014). β-Lactam antibiotics kill bacteria by inhibiting the transpeptidase reaction and preventing bacterial cell wall assembly (Prabhakaran et al., 1999). Many studies have recently shown that *V. alginolyticus* is highly resistant to ampicillin in environmental and aquatic isolates (Lajnef et al., 2012; Ripabelli et al., 2003). It is hypothesized that *V. alginolyticus* might already be naturally resistant to ampicillin. Notably, Clade IV isolates harbored significantly fewer AMR genes than strains from aquaculture environments or clinical settings, and isolates from wild hosts exhibited lower resistance than those from cultured hosts. The MAR index indicated that 32.61% of isolates were multidrug-resistant (MAR > 0.2), but high-level resistance (MAR = 0.6) was rare and restricted to a few host species (*A. djedaba* and *L. berbis*). These results strongly suggest that antibiotic use in intensive mariculture drives the emergence and spread of resistant *V. alginolyticus* strains, whereas wild fish-associated lineages retain a low-resistance phenotype due to limited anthropogenic selection pressure in Hong Kong coastal waters.

Virulence gene analysis revealed a conserved core repertoire that included *tlh*, T3SS1, and T6SS, but all isolates lacked the high-risk human-pathogenic determinants *tdh* and *ctxB*, indicating low zoonotic potential relative to pandemic *V. parahaemolyticus* and *V. cholerae*. Virulence loci were partitioned between the two chromosomes, with adhesion, motility, and secretion systems on Chromosome I and iron uptake and immune evasion functions on Chromosome II. Similar to AMR patterns, Clade IV and wild-fish isolates displayed a reduced virulence gene profile compared with aquaculture and pathogenic strains. We propose an evolutionary trade-off: in wild fish, *V. alginolyticus* maintains a low-virulence, commensal phenotype to ensure long-term host colonization; in aquaculture systems, high host density and stressful conditions favor strains with enhanced virulence to facilitate infection and transmission. This trade-off shapes the population structure and pathogenic potential of *V. alginolyticus* in contrasting marine ecosystems.

In prokaryotes, DNA methylation is critical not only for regulating defense systems but also for mediating environmental responses, including cell cycle control, gene expression, and virulence (Seong et al., 2021). A comprehensive understanding of DNA methylation is therefore essential for elucidating bacterial physiology. While bisulfite conversion-based methods have been the most popular method for detecting methylation, they are prone to false positives and expensive (Pajares et al., 2021). The emergence of third-generation sequencing, particularly nanopore technology, overcomes read-length limitations by providing ultra-long reads and enabling direct, genome-wide detection of modified bases (Chera et al., 2024). This is achieved by quantifying changes in the intensity of electrical currents as modified versus unmodified nucleotides pass through the nanopore (Liu et al., 2021). We generated complete circular genomes for 46 *V. alginolyticus* strains using third-generation nanopore sequencing and performed a comprehensive DNA methylome analysis. This study provides the first comprehensive methylomic characterization of *V. alginolyticus*, identifying 355 DNA methyltransferase genes and 140 methylation motifs dominated by 6mA modifications. A total of 71.90% of MTase genes were located in the accessory genome. Among these accessory-genomic MTases, 41.5% were harbored by mobile genetic elements (MGEs), with plasmids serving as the primary carrier, which indicates that MGEs act as major drivers for the diversification and horizontal dissemination of MTase genes in *V. alginolyticus*. Additionally, motif combinations were highly strain-specific and largely decoupled from phylogeny, which also indicates extensive epigenetic diversification. However, a conserved set of three motifs (“GATC”, “GNNGANNNNNNNNTTGG”, and “GGNAYNNNNNNNRTGG”) was exclusively shared by Clade IV strains, providing an epigenetic signature that supports the evolutionary coherence of this wild-fish-adapted clade. The strain-specific methylome observed here suggests that epigenetic heterogeneity contributes to niche adaptation and intraspecific diversification independently of genomic variation. These findings highlight the value of integrating methylomic data into bacterial population studies.

Some limitations should be considered. First, the sampling scope was geographically restricted to Hong Kong coastal waters and limited to a single seasonal sampling period (October and December 2022). Although the sampled wild fish belonged to 17 species, 8 genera, and 6 families and covered multiple trophic levels and vertical habitats, the strain population genomic and epigenomic characteristics observed herein may not fully represent the global or seasonal dynamic patterns of wild fish-derived *V. alginolyticus*. Therefore, long-term, cross-regional, and seasonal continuous sampling is required to verify the universality and temporal stability of the Clade IV low-virulence phenotype and its unique epigenetic signature. Second, the observed phylogenetic clustering of most wild fish-derived isolates within Clade IV may raise risks regarding potential clonal redundancy and overrepresentation of identical or highly similar strains in our dataset. To minimize sampling bias and eliminate redundant or cross-contaminated isolates, we implemented stringent procedures including individual packaging and processing of each fish and retention of only distinct genotypes from different hosts or unique variants from the same host. These steps ensured that the 46 isolates included in this study displayed appreciable genomic and epigenomic divergence, including variations in genome size, plasmid composition, and methylation profiles represented by 17 distinct motif combinations, most of which were strain-specific. Isolates within Clade IV were also obtained from eight distinct fish species, supporting their ecological and genetic distinctness. The observed phylogenetic distribution therefore likely reflects a common, well-adapted lineage endemic to the marine environment. Further expanded sampling across more locations and host species may help refine insights into the fine-scale population structure and evolutionary features of wild-fish-associated *V. alginolyticus*.

In summary, this study provides a bacterial genomic resource, nearly doubling the number of high-quality complete genomes available for *V. alginolyticus*, and establishes a unified pan-genomic and methylomic framework for understanding its evolution, host adaptation, virulence, and antimicrobial resistance. We identify a distinct, low-virulence Clade IV that is widely shared among wild marine fish, representing a specialized commensal lineage shaped by host ecology and minimal anthropogenic pressure. Virulence and resistance determinants are primarily chromosomal, with plasmids contributing to environmental adaptation rather than pathogenicity. The strain-specific epigenome further underscores the complexity of *V. alginolyticus* adaptation in marine ecosystems. These findings provide critical insights for One Health surveillance, mariculture management, and risk assessment of *V. alginolyticus* in coastal environments.

## 5 Conclusion

This study establishes a comprehensive pan-genomic and methylomic framework for *V. alginolyticus* and reveals a phylogenetically distinct Clade IV adapted to wild marine fish, characterized by low virulence and unique methylome. We demonstrate that resistance and virulence genes are predominantly chromosomal in these bacteria, while plasmids support environmental adaptation. The methylome is highly strain-specific and provides an epigenetic signature for Clade IV. Our results expand genomic resources for this important pathogen and inform One Health surveillance and risk assessment in marine ecosystems.

## Acknowledgments

This research was funded by the APRC-CityU New Research Initiatives/Infrastructure Support (9610574) and the SIRG-CityU Strategic Interdisciplinary Research Grant (7020090). This work was also supported by the Innovation and Technology Commission (ITC) of the Hong Kong SAR Government (9448002), which provides regular research funding support to the State Key Laboratory of Marine Environmental Health. However, any opinions, findings, conclusions, or recommendations expressed in this publication do not reflect the views of the Hong Kong SAR Government nor the ITC. We thank the fishermen for assistance with field sampling in Tolo Harbor, Hong Kong.

## Data availability

All raw sequencing reads and complete genome assemblies generated in this study have been deposited in the NCBI Sequence Read Archive (SRA) and GenBank under BioProject accession number PRJNA1466115. Public genomes used in this study are available from the NCBI RefSeq database under the accession numbers listed in Supplementary Table 3. All supporting data, including phylogenetic trees, gene annotations, and methylation motifs, are available in the supplementary materials and from the corresponding author upon reasonable request.

## Authors’ contributions

Z.L. and Y.Z. performed sample collection, bacterial isolation, genome sequencing, and bioinformatic analysis. M.Y. and R.L. contributed to phylogenetic and pan-genomic analysis. W.C. conceived and supervised the project, acquired funding, and designed the experiments. Z.L., Y.Z., and W.C. wrote the manuscript with input from all authors. All authors read and approved the final manuscript.

## References

Alcock, B.P., Huynh, W., Chalil, R., Smith, K.W., Raphenya, Amogelang R., Wlodarski, M.A., Edalatmand, A., Petkau, A., Syed, S.A., Tsang, K.K., Baker, S.J.C., Dave, M., McCarthy, Madeline C., Mukiri, K.M., Nasir, J.A., Golbon, B., Imtiaz, H., Jiang, X., Kaur, K., Kwong, M., Liang, Z.C., Niu, K.C., Shan, P., Yang, J.Y.J., Gray, Kristen L., Hoad, G.R., Jia, B., Bhando, T., Carfrae, Lindsey A., Farha, Maya A., French, S., Gordzevich, R., Rachwalski, K., Tu, Megan M., Bordeleau, E., Dooley, D., Griffiths, E., Zubyk, H.L., Brown, E.D., Maguire, F., Beiko, Robert G., Hsiao, W.W.L., Brinkman, F.S.L., Van Domselaar, G., McArthur, A.G., 2023. CARD 2023: expanded curation, support for machine learning, and resistome prediction at the Comprehensive Antibiotic Resistance Database. Nucleic Acids Research 51(D1), D690–D699. 10.1093/nar/gkac920

Altekruse, S.F., Bishop, R.D., Baldy, L.M., Thompson, S.G., Wilson, S.A., Ray, B.J., Griffin, P.M., 2000. Vibrio gastroenteritis in the US Gulf of Mexico region: the role of raw oysters. Epidemiol Infect 124(3), 489–495. 10.1017/s0950268899003714

Arndt, D., Grant, J.R., Marcu, A., Sajed, T., Pon, A., Liang, Y., Wishart, D.S., 2016. PHASTER: a better, faster version of the PHAST phage search tool. Nucleic Acids Research 44(W1), W16–W21. 10.1093/nar/gkw387

Bailey, T.L., Johnson, J., Grant, C.E., Noble, W.S., 2015. The MEME Suite. Nucleic Acids Research 43(W1), W39–W49. 10.1093/nar/gkv416

Balaban, M., Jiang, Y., Zhu, Q., McDonald, D., Knight, R., Mirarab, S., 2024. Generation of accurate, expandable phylogenomic trees with uDance. Nature Biotechnology 42(5), 768–777. 10.1038/s41587-023-01868-8

Beaulaurier, J., Schadt, E.E., Fang, G., 2019. Deciphering bacterial epigenomes using modern sequencing technologies. Nat Rev Genet 20(3), 157–172. 10.1038/s41576-018-0081-3

Bertelli, C., Brinkman, F.S.L., 2018. Improved genomic island predictions with IslandPath-DIMOB. Bioinformatics 34(13), 2161–2167. 10.1093/bioinformatics/bty095

Camacho, C., Coulouris, G., Avagyan, V., Ma, N., Papadopoulos, J., Bealer, K., Madden, T.L., 2009. BLAST+: architecture and applications. BMC Bioinformatics 10(1), 421. 10.1186/1471-2105-10-421

Cao, J., Zhang, J., Ma, L., Li, L., Zhang, W., Li, J., 2018. Identification of fish source Vibrio alginolyticus and evaluation of its bacterial ghosts vaccine immune effects. MicrobiologyOpen 7(3), e00576. 10.1002/mbo3.576

Casadesús, J., 2016. Bacterial DNA Methylation and Methylomes. Adv Exp Med Biol 945, 35–61. 10.1007/978-3-319-43624-1_3

CDC, C.f.D.C.a.P., 2012. Cholera and Other Vibrio Illness Surveillance (COVIS), summary data, 2008–2012. Atlanta, GA: US Department of Health and Human Services. https://www.cdc.gov/vibrio/surveillance.html Accessed 31 July 2020.

Ceccarelli, D., Amaro, C., Romalde, J.L., Suffredini, E., Vezzulli, L., 2019. Vibrio Species, in: Food Microbiology. pp. 347–388.

Chao, M.C., Zhu, S., Kimura, S., Davis, B.M., Schadt, E.E., Fang, G., Waldor, M.K., 2015. A cytosine methytransferase modulates the cell envelope stress response in the cholera pathogen. PLoS genetics 11(11), e1005666.

Chera, A., Stancu-Cretu, M., Zabet, N.R., Bucur, O., 2024. Shedding light on DNA methylation and its clinical implications: the impact of long-read-based nanopore technology. Epigenetics & Chromatin 17(1), 39. 10.1186/s13072-024-00558-2

CLSI, 2018. Performance Standards for Antimicrobial Disk Susceptibility Tests. 13th ed. CLSI standard M02 Wayne, PA: Clinical and Laboratory Standards Institute; 2018.

de Souza Valente, C., Wan, A.H.L., 2021. Vibrio and major commercially important vibriosis diseases in decapod crustaceans. Journal of invertebrate pathology 181, 107527. 10.1016/j.jip.2020.107527

Deng, Y., Xu, L., Chen, H., Liu, S., Guo, Z., Cheng, C., Ma, H., Feng, J., 2020. Prevalence, virulence genes, and antimicrobial resistance of Vibrio species isolated from diseased marine fish in South China. Sci Rep 10(1), 14329. 10.1038/s41598-020-71288-0

Durand, G.A., Raoult, D., Dubourg, G., 2019. Antibiotic discovery: history, methods and perspectives. International journal of antimicrobial agents 53(4), 371–382. 10.1016/j.ijantimicag.2018.11.010

Edgar, R.C., 2022. Muscle5: High-accuracy alignment ensembles enable unbiased assessments of sequence homology and phylogeny. Nature Communications 13(1), 6968. 10.1038/s41467-022-34630-w

Elmahdi, S., DaSilva, L.V., Parveen, S., 2016. Antibiotic resistance of Vibrio parahaemolyticus and Vibrio vulnificus in various countries: A review. Food microbiology 57, 128–134. 10.1016/j.fm.2016.02.008

Emms, D.M., Kelly, S., 2019. OrthoFinder: phylogenetic orthology inference for comparative genomics. Genome Biology 20(1), 238. 10.1186/s13059-019-1832-y

He, F., Chen, S., Zhang, Y., Chai, K., Zhang, Q., Kong, W., Qu, S., Chen, L., Zhang, F., Li, M., Wang, X., Lv, H., Zhang, T., He, X., Li, X., Li, Y., Li, X., Jiang, X., Xu, M., Sod, B., Kang, J., Zhang, X., Long, R., Yang, Q., 2025. Pan-genomic analysis highlights genes associated with agronomic traits and enhances genomics-assisted breeding in alfalfa. Nature Genetics 57(5), 1262–1273. 10.1038/s41588-025-02164-8

Hlady, W.G., Klontz, K.C., 1996. The epidemiology of Vibrio infections in Florida, 1981-1993. J Infect Dis 173(5), 1176–1183. 10.1093/infdis/173.5.1176

Huang, Z., Li, Y., Yu, K., Ma, L., Pang, B., Qin, Q., Li, J., Wang, D., Gao, H., Kan, B., 2024. Genome-wide expanding of genetic evolution and potential pathogenicity in Vibrio alginolyticus. Emerg Microbes Infect 13(1), 2350164. 10.1080/22221751.2024.2350164

Hyatt, D., Chen, G.-L., LoCascio, P.F., Land, M.L., Larimer, F.W., Hauser, L.J., 2010. Prodigal: prokaryotic gene recognition and translation initiation site identification. BMC Bioinformatics 11(1), 119. 10.1186/1471-2105-11-119

Igbinosa, E.O., 2016. Detection and Antimicrobial Resistance of Vibrio Isolates in Aquaculture Environments: Implications for Public Health. Microbial drug resistance (Larchmont, N.Y.) 22(3), 238–245. 10.1089/mdr.2015.0169

Jacobs Slifka, K.M., Newton, A.E., Mahon, B.E., 2017. Vibrio alginolyticus infections in the USA, 1988-2012. Epidemiol Infect 145(7), 1491–1499. 10.1017/s0950268817000140

Krumperman, P.H., 1983. Multiple antibiotic resistance indexing of Escherichia coli to identify high-risk sources of fecal contamination of foods. Applied and environmental microbiology 46(1), 165–170. 10.1128/aem.46.1.165-170.1983

Lajnef, R., Snoussi, M., Romalde, J.L., Nozha, C., Hassen, A., 2012. Comparative study on the antibiotic susceptibility and plasmid profiles of Vibrio alginolyticus strains isolated from four Tunisian marine biotopes. World journal of microbiology & biotechnology 28(12), 3345–3363. 10.1007/s11274-012-1147-6

Lassalle, F., Al-Shalali, S., Al-Hakimi, M., Njamkepo, E., Bashir, I.M., Dorman, M.J., Rauzier, J., Blackwell, G.A., Taylor-Brown, A., Beale, M.A., Cazares, A., Al-Somainy, A.A., Al-Mahbashi, A., Almoayed, K., Aldawla, M., Al-Harazi, A., Quilici, M.-L., Weill, F.-X., Dhabaan, G., Thomson, N.R., 2023. Genomic epidemiology reveals multidrug resistant plasmid spread between Vibrio cholerae lineages in Yemen. Nature Microbiology 8(10), 1787–1798. 10.1038/s41564-023-01472-1

Liu, B., Zheng, D., Jin, Q., Chen, L., Yang, J., 2019. VFDB 2019: a comparative pathogenomic platform with an interactive web interface. Nucleic Acids Research 47(D1), D687–D692. 10.1093/nar/gky1080

Liu, C.H., Cheng, W., Hsu, J.P., Chen, J.C., 2004. Vibrio alginolyticus infection in the white shrimp Litopenaeus vannamei confirmed by polymerase chain reaction and 16S rDNA sequencing. Dis Aquat Organ 61(1-2), 169–174. 10.3354/dao061169

Liu, M., Li, X., Xie, Y., Bi, D., Sun, J., Li, J., Tai, C., Deng, Z., Ou, H.-Y., 2019. ICEberg 2.0: an updated database of bacterial integrative and conjugative elements. Nucleic Acids Research 47(D1), D660–D665. 10.1093/nar/gky1123

Liu, Y., Rosikiewicz, W., Pan, Z., Jillette, N., Wang, P., Taghbalout, A., Foox, J., Mason, C., Carroll, M., Cheng, A., Li, S., 2021. DNA methylation-calling tools for Oxford Nanopore sequencing: a survey and human epigenome-wide evaluation. Genome Biology 22(1), 295. 10.1186/s13059-021-02510-z

Lu, B., Guo, Z., Liu, X., Ni, Y., Xu, L., Huang, J., Li, T., Feng, T., Li, R., Deng, X., 2025. Comprehensive comparison of the third-generation sequencing tools for bacterial 6mA profiling. Nature Communications 16(1), 3982. 10.1038/s41467-025-59187-2

Ma, Y.-x., Wang, X.-d., Li, X.-m., 2025. The emerging role of DNA methylation in the pathogenicity of bacterial pathogens. Journal of Bacteriology 207(8), e00108–00125. doi:10.1128/jb.00108-25

Manni, M., Berkeley, M.R., Seppey, M., Simão, F.A., Zdobnov, E.M., 2021. BUSCO Update: Novel and Streamlined Workflows along with Broader and Deeper Phylogenetic Coverage for Scoring of Eukaryotic, Prokaryotic, and Viral Genomes. Molecular Biology and Evolution 38(10), 4647–4654. 10.1093/molbev/msab199

Martin, S., Fournes, F., Ambrosini, G., Iseli, C., Bojkowska, K., Marquis, J., Guex, N., Collier, J., 2024. DNA methylation by CcrM contributes to genome maintenance in the Agrobacterium tumefaciens plant pathogen. Nucleic Acids Research 52(19), 11519–11535. 10.1093/nar/gkae757

Mitchell, S.M., Ullman, J.L., Teel, A.L., Watts, R.J., 2014. pH and temperature effects on the hydrolysis of three β-lactam antibiotics: ampicillin, cefalotin and cefoxitin. Sci Total Environ 466-467, 547–555. 10.1016/j.scitotenv.2013.06.027

Mohamad, N., Mohd Roseli, F.A., Azmai, M.N.A., Saad, M.Z., Md Yasin, I.S., Zulkiply, N.A., Nasruddin, N.S., 2019. Natural Concurrent Infection of Vibrio harveyi and V. alginolyticus in Cultured Hybrid Groupers in Malaysia. J Aquat Anim Health 31(1), 88–96. 10.1002/aah.10055

Muzzi, A., Masignani, V., Rappuoli, R., 2007. The pan-genome: towards a knowledge-based discovery of novel targets for vaccines and antibacterials. Drug Discovery Today 12(11), 429–439. 10.1016/j.drudis.2007.04.008

Nathamuni, S., Jangam, A.K., Katneni, V.K., Selvaraj, A., Krishnan, K., Kumar, S., Avunje, S., Balasubramaniam, S., Grover, M., Alavandi, S.V., Koyadan, V.K., 2019. Insights on genomic diversity of Vibrio spp. through Pan-genome analysis. Annals of Microbiology 69(13), 1547–1555. 10.1007/s13213-019-01539-7

Oliveira, P.H., Fang, G., 2021. Conserved DNA Methyltransferases: A Window into Fundamental Mechanisms of Epigenetic Regulation in Bacteria. Trends Microbiol 29(1), 28–40. 10.1016/j.tim.2020.04.007

Pajares, M.J., Palanca-Ballester, C., Urtasun, R., Alemany-Cosme, E., Lahoz, A., Sandoval, J., 2021. Methods for analysis of specific DNA methylation status. Methods 187, 3–12. 10.1016/j.ymeth.2020.06.021

Prabhakaran, K., Harris, E.B., Randhawa, B., 1999. Bactericidal action of ampicillin/sulbactam against intracellular mycobacteria. International journal of antimicrobial agents 13(2), 133–135. 10.1016/s0924-8579(99)00101-6

Quinlan, A.R., Hall, I.M., 2010. BEDTools: a flexible suite of utilities for comparing genomic features. Bioinformatics 26(6), 841–842. 10.1093/bioinformatics/btq033

Rameshkumar, P., Nazar, A.K.A., Pradeep, M.A., Kalidas, C., Jayakumar, R., Tamilmani, G., Sakthivel, M., Samal, A.K., Sirajudeen, S., Venkatesan, V., Nazeera, B.M., 2017. Isolation and characterization of pathogenic Vibrio alginolyticus from sea cage cultured cobia (Rachycentron canadum (Linnaeus 1766)) in India. Lett Appl Microbiol 65(5), 423–430. 10.1111/lam.12800

Riggio, M.P., Lappin, D.F., Bennett, D., 2014. Bacteria and Toll-like receptor and cytokine mRNA expression profiles associated with canine arthritis. Vet Immunol Immunopathol 160(3-4), 158–166. 10.1016/j.vetimm.2014.04.004

Ripabelli, G., Sammarco, M.L., McLauchlin, J., Fanelli, I., 2003. Molecular characterisation and antimicrobial resistance of Vibrio vulnificus and Vibrio alginolyticus isolated from mussels (Mytilus galloprovincialis). Systematic and applied microbiology 26(1), 119–126. 10.1078/072320203322337407

Roberts, R.J., Vincze, T., Posfai, J., Macelis, D., 2003. REBASE: restriction enzymes and methyltransferases. Nucleic Acids Research 31(1), 418–420. 10.1093/nar/gkg069

Rodríguez-Beltrán, J., DelaFuente, J., León-Sampedro, R., MacLean, R.C., San Millán, Á., 2021. Beyond horizontal gene transfer: the role of plasmids in bacterial evolution. Nature Reviews Microbiology 19(6), 347–359. 10.1038/s41579-020-00497-1

Seemann, T., 2014. Prokka: rapid prokaryotic genome annotation. Bioinformatics 30(14), 2068–2069. 10.1093/bioinformatics/btu153

Seong, H.J., Han, S.W., Sul, W.J., 2021. Prokaryotic DNA methylation and its functional roles. J Microbiol 59(3), 242–248. 10.1007/s12275-021-0674-y

Sganga, G., Cozza, V., Spanu, T., Spada, P.L., Fadda, G., 2009. Global climate change and wound care: case study of an off-season vibrio alginolyticus infection in a healthy man. Ostomy Wound Manage 55(4), 60–62.

Snipen, L., Liland, K.H., 2015. micropan: an R-package for microbial pan-genomics. BMC Bioinformatics 16, 79. 10.1186/s12859-015-0517-0

Stamatakis, A., 2014. RAxML version 8: a tool for phylogenetic analysis and post-analysis of large phylogenies. Bioinformatics 30(9), 1312–1313. 10.1093/bioinformatics/btu033

Tettelin, H., Masignani, V., Cieslewicz, M.J., Donati, C., Medini, D., Ward, N.L., Angiuoli, S.V., Crabtree, J., Jones, A.L., Durkin, A.S., DeBoy, R.T., Davidsen, T.M., Mora, M., Scarselli, M., Margarit y Ros, I., Peterson, J.D., Hauser, C.R., Sundaram, J.P., Nelson, W.C., Madupu, R., Brinkac, L.M., Dodson, R.J., Rosovitz, M.J., Sullivan, S.A., Daugherty, S.C., Haft, D.H., Selengut, J., Gwinn, M.L., Zhou, L., Zafar, N., Khouri, H., Radune, D., Dimitrov, G., Watkins, K., O’Connor, K.J.B., Smith, S., Utterback, T.R., White, O., Rubens, C.E., Grandi, G., Madoff, L.C., Kasper, D.L., Telford, J.L., Wessels, M.R., Rappuoli, R., Fraser, C.M., 2005. Genome analysis of multiple pathogenic isolates of *Streptococcus agalactiae*: Implications for the microbial &#x201c;pan-genome&#x201d. Proceedings of the National Academy of Sciences 102(39), 13950–13955. doi:10.1073/pnas.0506758102

Tisza, M.J., Smith, D.D.N., Clark, A.E., Youn, J.H., Khil, P.P., Dekker, J.P., 2023. Roving methyltransferases generate a mosaic epigenetic landscape and influence evolution in Bacteroides fragilis group. Nat Commun 14(1), 4082. 10.1038/s41467-023-39892-6

Uh, Y., Park, J.S., Hwang, G.Y., Jang, I.H., Yoon, K.J., Park, H.C., Hwang, S.O., 2001. Vibrio alginolyticus acute gastroenteritis: report of two cases. Clin Microbiol Infect 7(2), 104–106. 10.1046/j.1469-0691.2001.00207.x

Van Goethem, M.W., Pierneef, R., Bezuidt, O.K.I., Van De Peer, Y., Cowan, D.A., Makhalanyane, T.P., 2018. A reservoir of ’historical’ antibiotic resistance genes in remote pristine Antarctic soils. Microbiome 6(1), 40. 10.1186/s40168-018-0424-5

Wang, K., Zhang, C., Munang’andu, H.M., Xu, C., Cai, W., Yan, X., Tao, Z., 2025. Comparative Genomic Analysis of Two Vibrio harveyi Strains from Larimichthys crocea with Divergent Virulence Profiles. Microorganisms 13(5), 1129.

Wang, T., Yao, L., Qu, M., Wang, L., Li, F., Tan, Z., Wang, P., Jiang, Y., 2022. Whole genome sequencing and antimicrobial resistance analysis of Vibrio parahaemolyticus Vp2015094 carrying an antimicrobial-resistant plasmid. Journal of Global Antimicrobial Resistance 30, 47–49. 10.1016/j.jgar.2022.05.025

Wang, Z., Wang, B., Chen, G., Jian, J., Lu, Y., Xu, Y., Wu, Z., 2016. Transcriptome analysis of the pearl oyster (Pinctada fucata) hemocytes in response to Vibrio alginolyticus infection. Gene 575(2 Pt 2), 421–428. 10.1016/j.gene.2015.09.014

Xie, Z., Tang, H., 2017. ISEScan: automated identification of insertion sequence elements in prokaryotic genomes. Bioinformatics 33(21), 3340–3347. 10.1093/bioinformatics/btx433

Yang, B., Zhai, S., Li, X., Tian, J., Li, Q., Shan, H., Liu, S., 2021. Identification of Vibrio alginolyticus as a causative pathogen associated with mass summer mortality of the Pacific Oyster (Crassostrea gigas) in China. Aquaculture 535, 736363. 10.1016/j.aquaculture.2021.736363

Yin, Z., Liang, J., Zhang, M., Chen, B., Yu, Z., Tian, X., Deng, X., Peng, L., 2024. Pan-genome insights into adaptive evolution of bacterial symbionts in mixed host-microbe symbioses represented by human gut microbiota Bacteroides cellulosilyticus. Science of The Total Environment 927, 172251. 10.1016/j.scitotenv.2024.172251

Yu, G., Wang, L.-G., Han, Y., He, Q.-Y., 2012. clusterProfiler: an R Package for Comparing Biological Themes Among Gene Clusters. OMICS: A Journal of Integrative Biology 16(5), 284–287. 10.1089/omi.2011.0118

Yu, Y., Li, H., Wang, Y., Zhang, Z., Liao, M., Rong, X., Li, B., Wang, C., Ge, J., Zhang, X., 2022. Antibiotic resistance, virulence and genetic characteristics of Vibrio alginolyticus isolates from aquatic environment in costal mariculture areas in China. Mar Pollut Bull 185(Pt A), 114219. 10.1016/j.marpolbul.2022.114219

Zhang, P., Liang, J., Mai, W., Wu, Y., Dai, J., Wei, Y., 2021. The efficiency of integrated wastewater treatment plant for pollutant removal from industrial-scale lincomycin production. Journal of Water Process Engineering 42, 102133. 10.1016/j.jwpe.2021.102133

Zhang, X.H., Austin, B., 2005. Haemolysins in Vibrio species. Journal of Applied Microbiology 98(5), 1011–1019. 10.1111/j.1365-2672.2005.02583.x

Zhang, Y., Shao, Y., Gao, S., Li, R., Cai, W., 2026. Prolonged starvation drives epigenetic remodeling: Insights from DNA methylation profiling in the aquatic pathogen Flavobacterium columnare. Water Biology and Security, 100604. 10.1016/j.watbs.2026.100604

