## Supplementary material for "A pan-genomic and methylomic analysis reveals a distinct signature in *Vibrio alginolyticus* isolated from wild fish": Figure S1-Figure S12

**Supplementary materials**

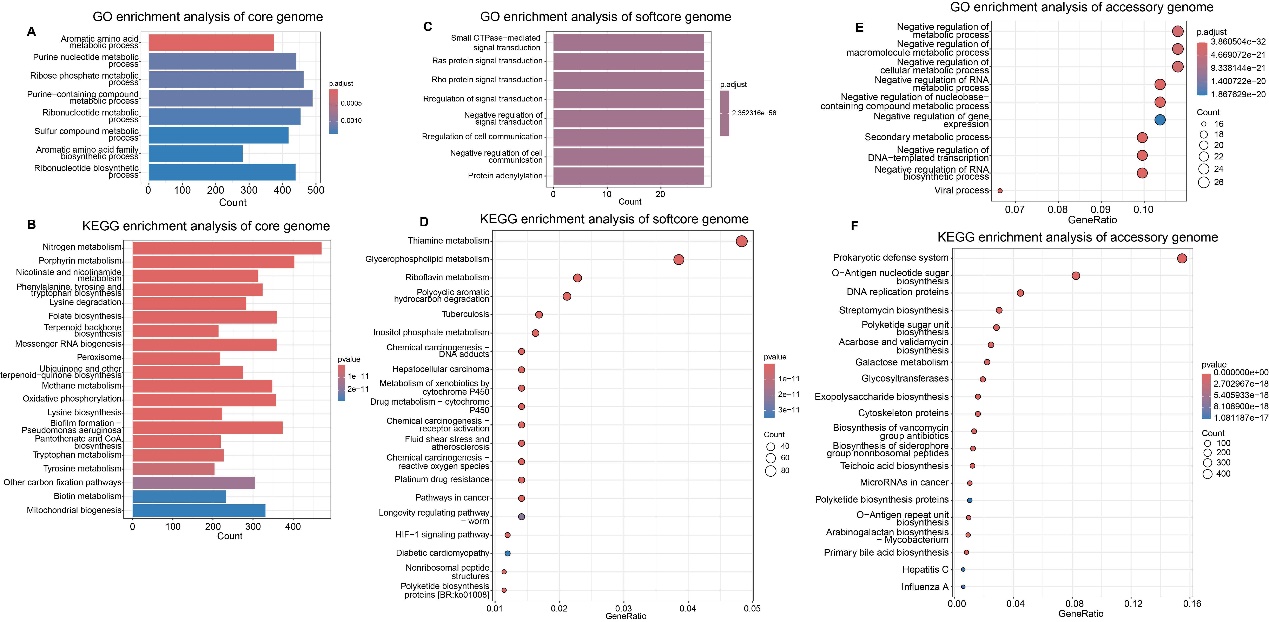

Figure S1. GO and KEGG enrichment analysis of core and accessory genomes.

A-B. GO and KEGG enrichment analysis of the core genome, respectively. C-D. GO and KEGG enrichment analysis of soft-core genome, respectively. E-F. GO and KEGG enrichment analysis of the accessory genome, respectively.

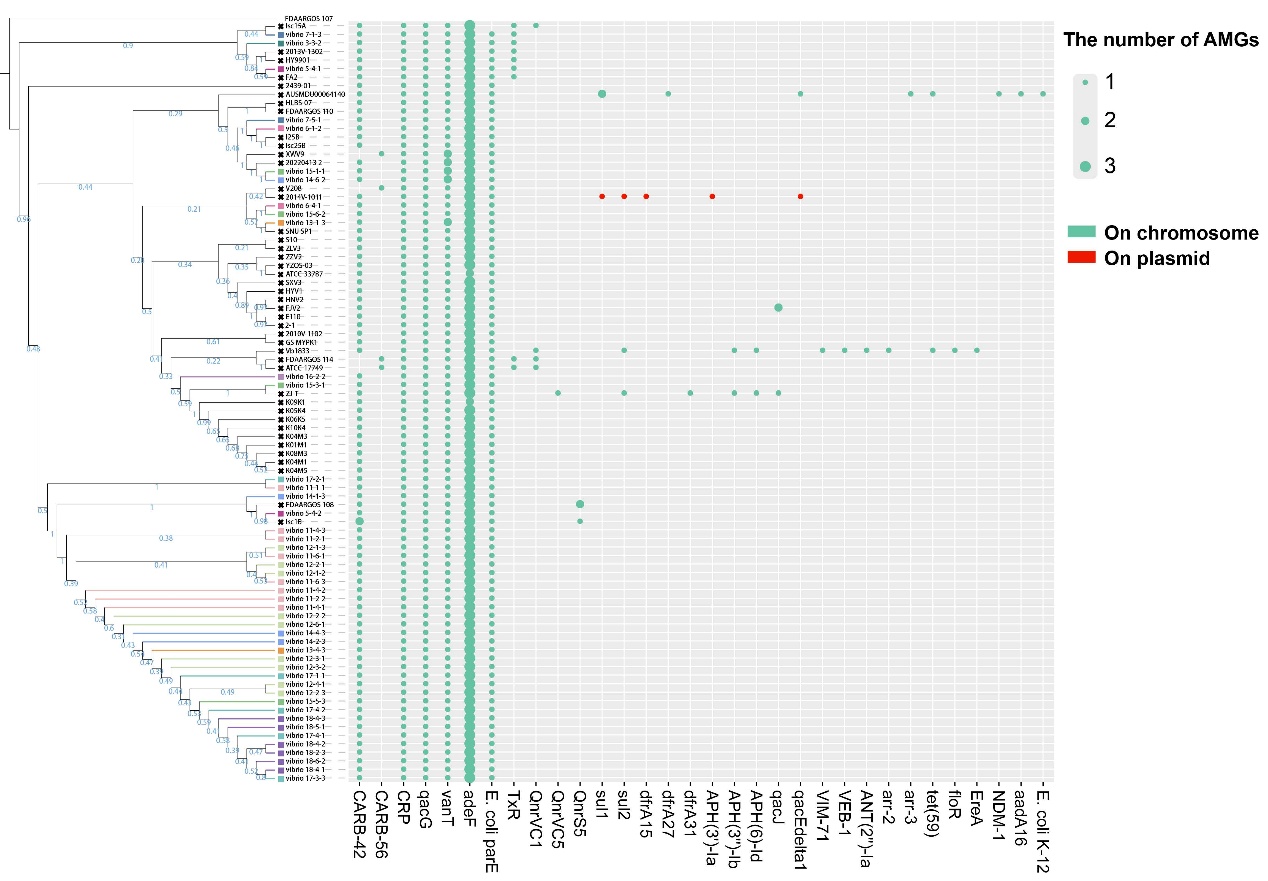

Figure S2. The distribution of antimicrobial resistance (AMR) genes in the 89 *V. alginolyticus* strains. The different colors represent the different locations of AMR genes in the genome of *V. alginolyticus* (The green color shows the AMR genes located on the chromosome, and the red color shows the AMR genes located on the plasmid).

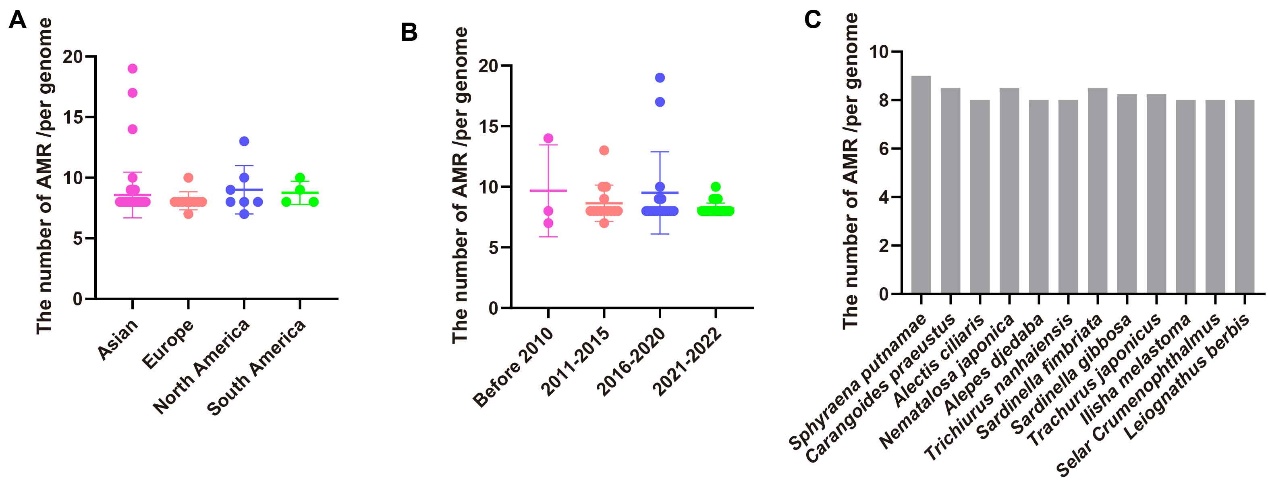

Figure S3. The relationship between the number of AMR genes in *V. alginolyticus* strains and isolated location (A), time (B), and fish species (C).

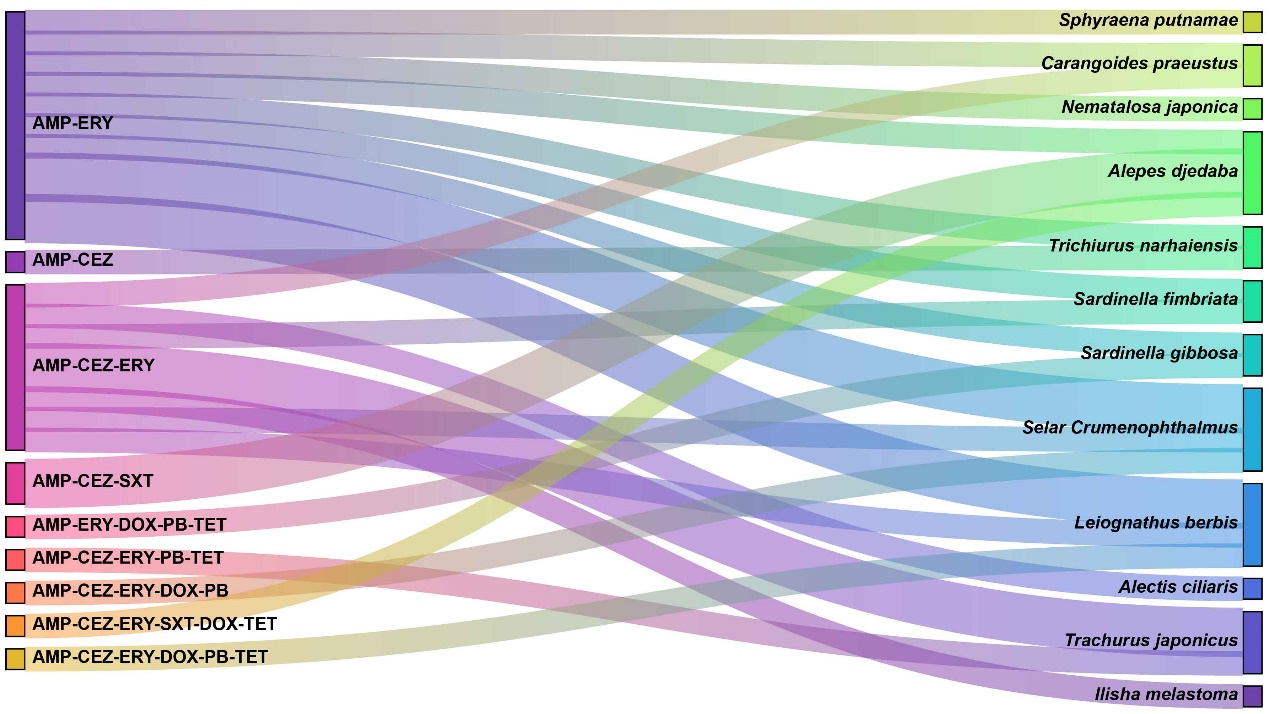

Figure S4. The multiple antibiotic resistance patterns analysis of *V. alginolyticus* isolates. The left side showed the multiple antibiotic resistance phenotypes, the right side showed the fish species to which the isolates belonged, and the width of the middle rectangular bar represented the number of resistant isolates.

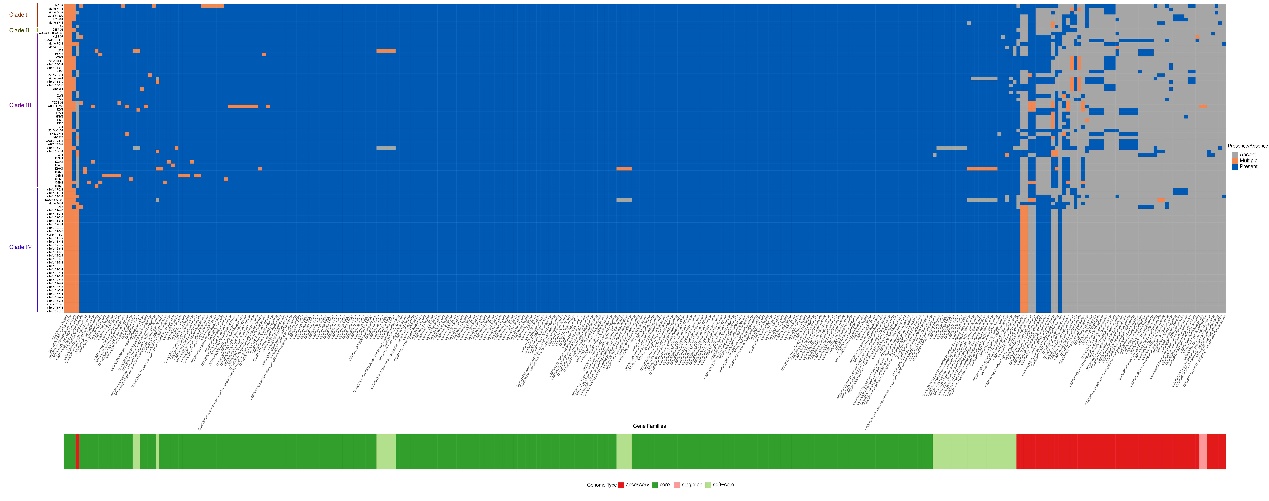

Figure S5. The distribution of the virulence factor genes in the core and accessory genomes.

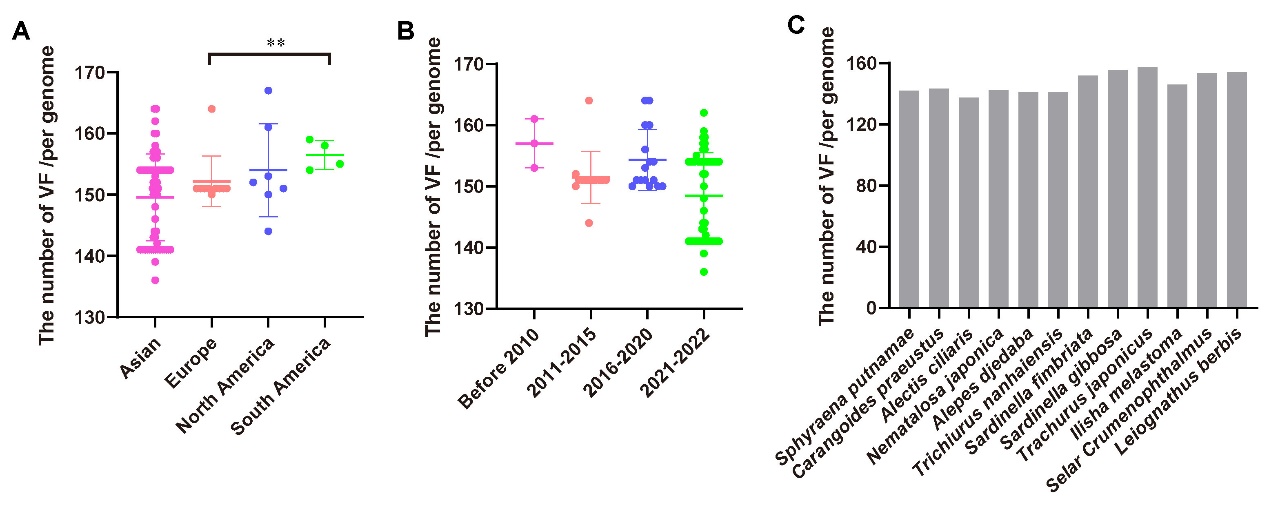

Figure S6. The relationship between the number of virulence genes in *V. alginolyticus* strains and isolated location (A), time (B), and fish species (C).

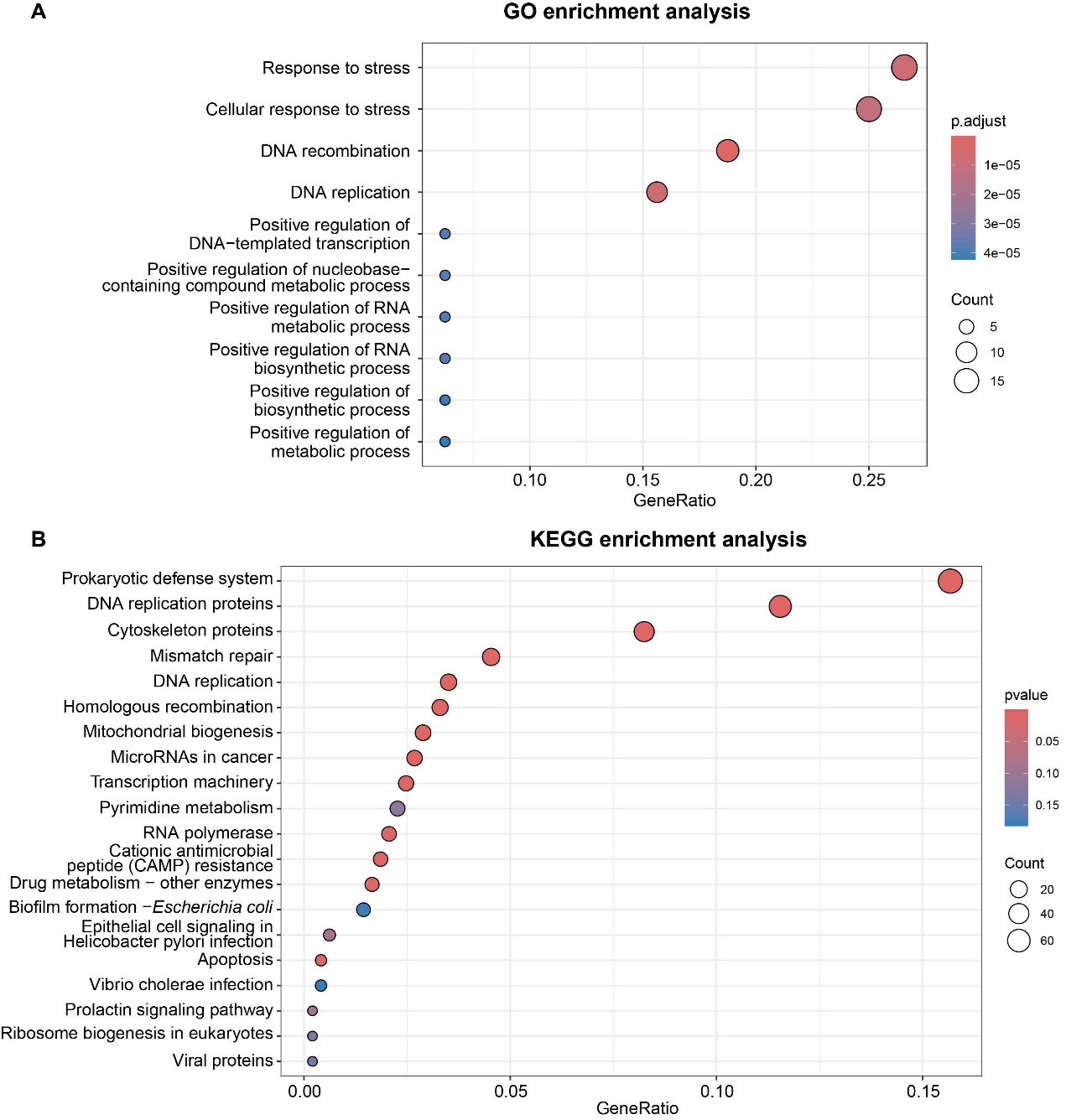

Figure S7. A and B. GO and KEGG enrichment analysis of genes in the plasmids, respectively.

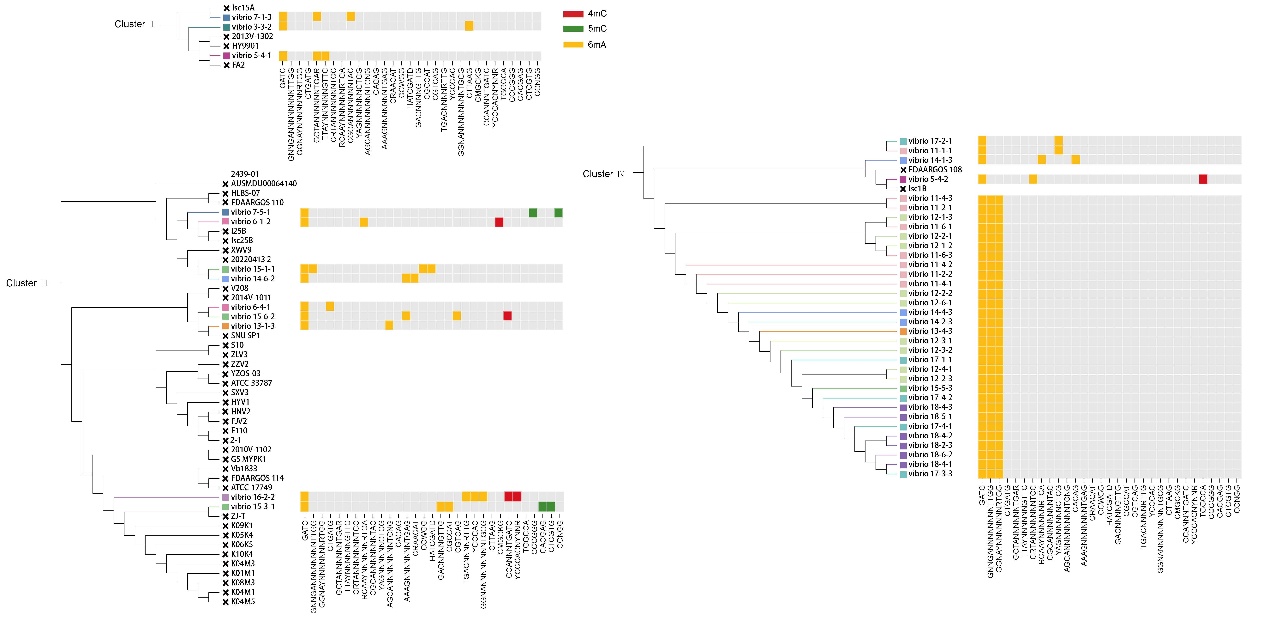

Figure S8. The distribution of methylation motifs across the 46 *V. alginolyticus* genomes isolated in this study and their correspondence with the phylogenetic tree

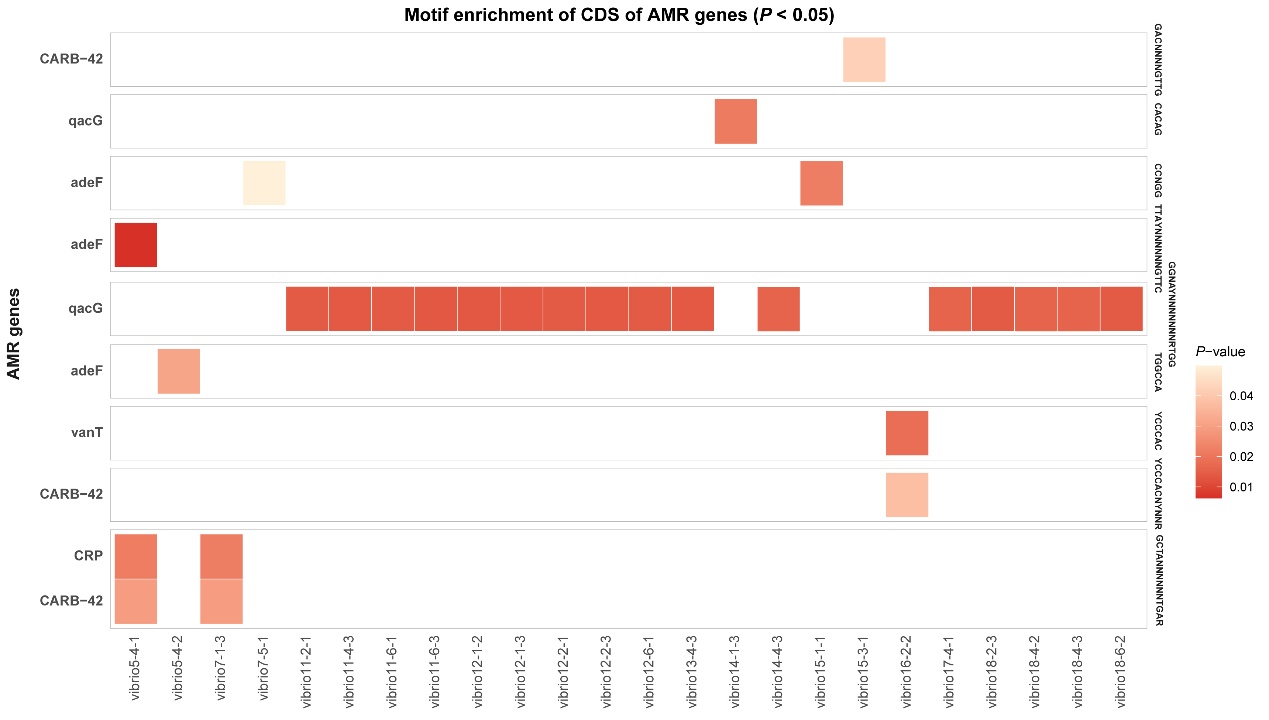

Figure S9. Distribution of significant motif enrichment in the coding regions of AMR genes.

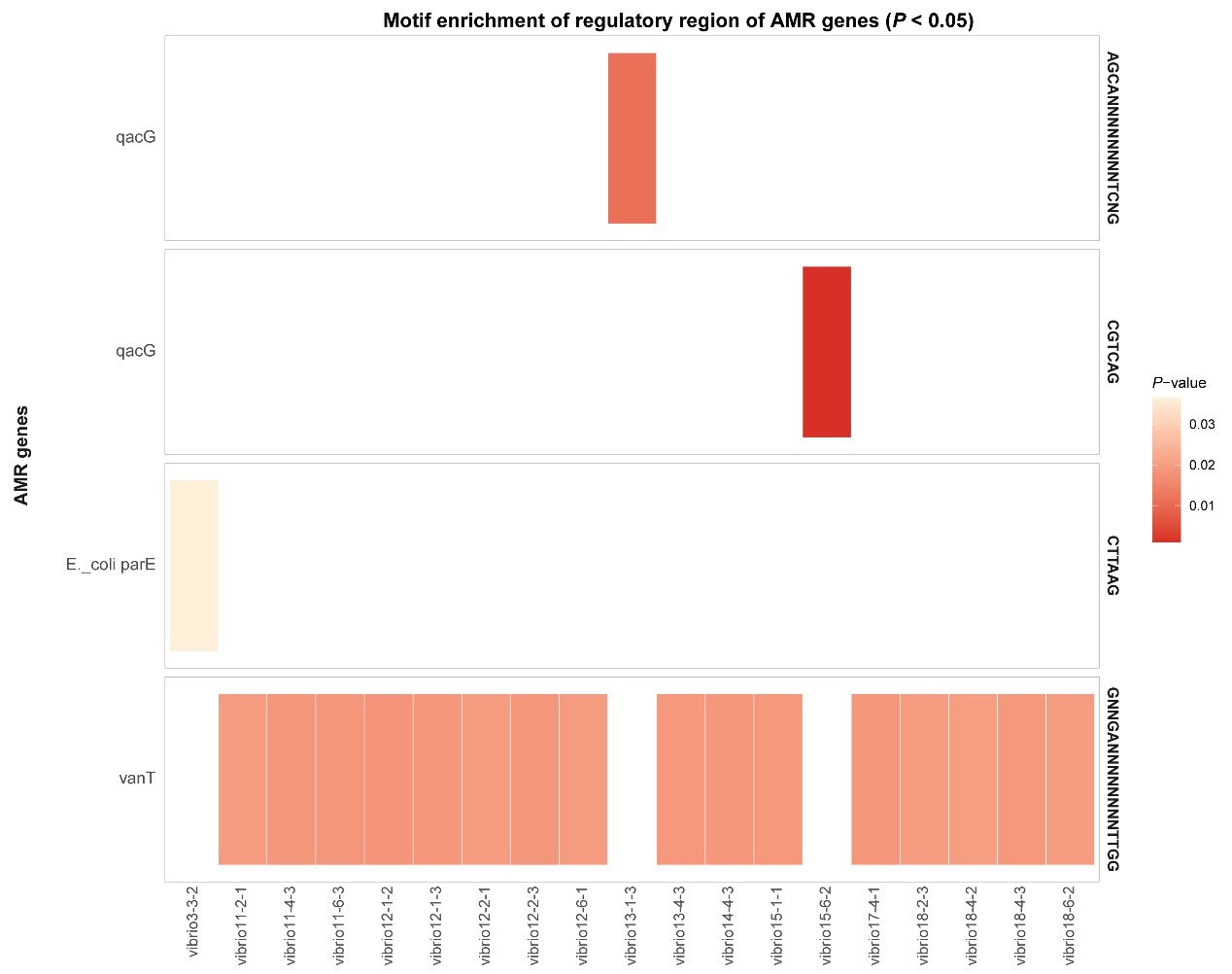

Figure S10. Distribution of significant motif enrichment in the regulatory regions of AMR genes.

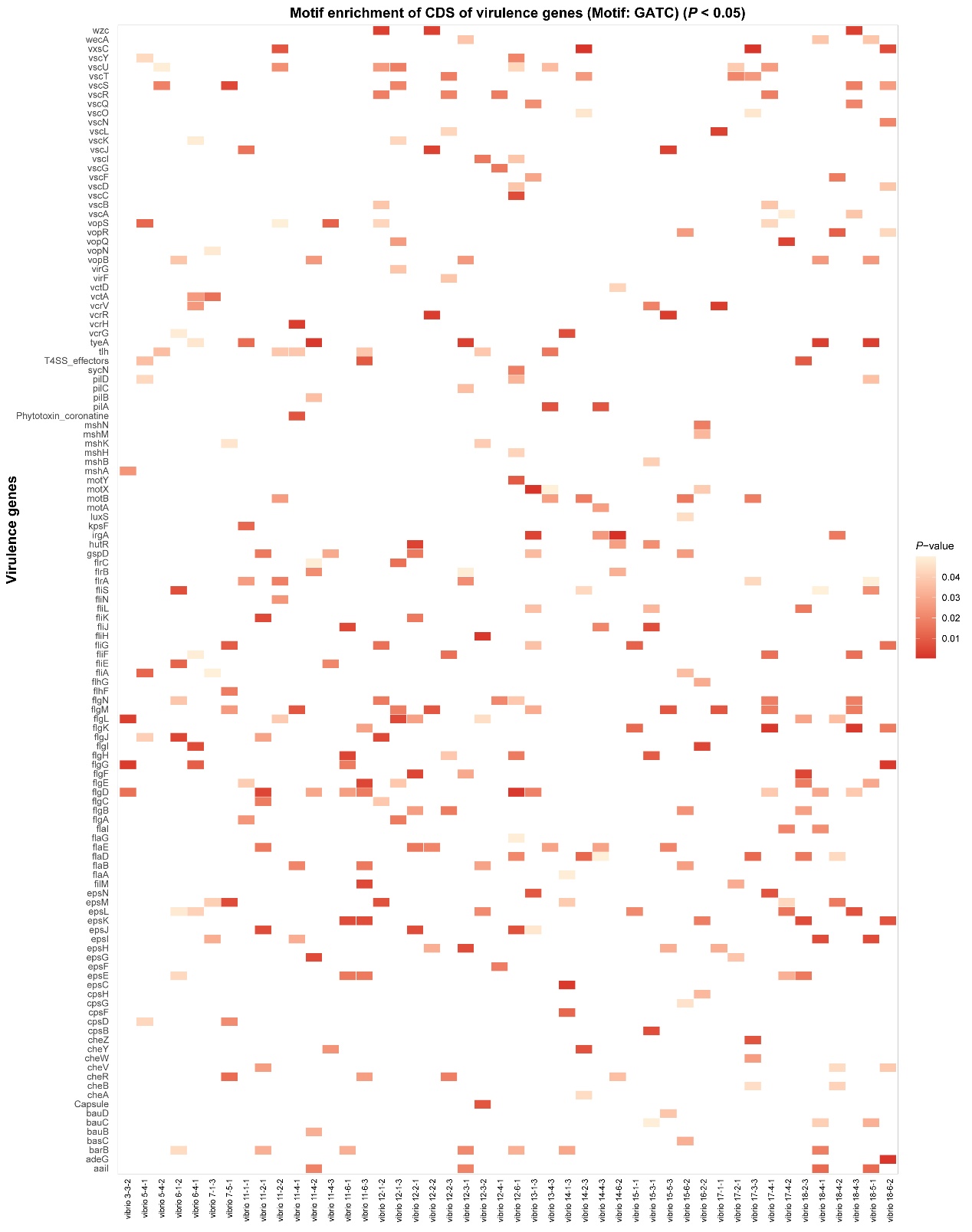

Figure S11. GATC motif enrichment in the coding regions of virulence genes.

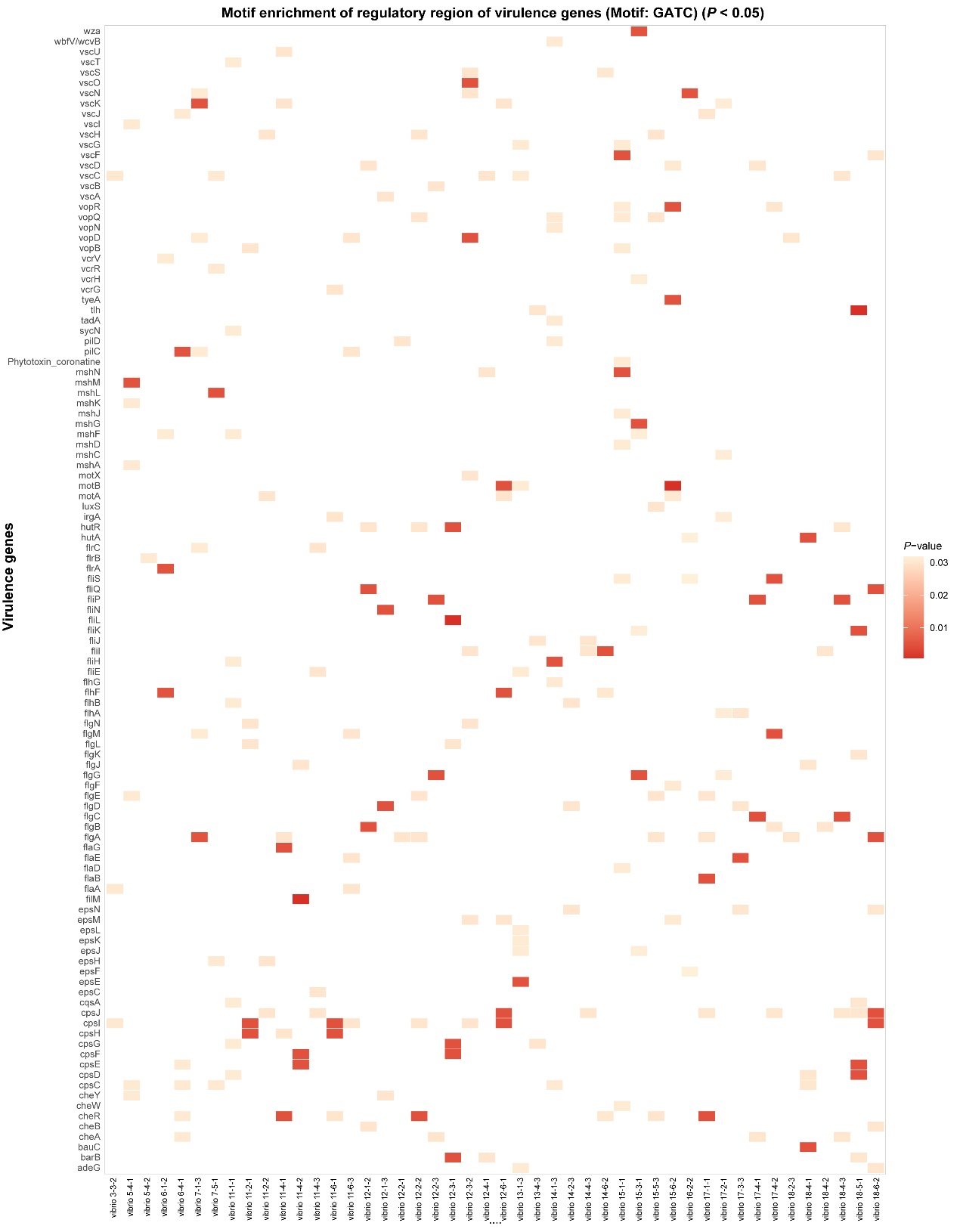

Figure S12. GATC motif enrichment in the regulatory regions of virulence genes.
